# Controlling seizure propagation in large-scale brain networks

**DOI:** 10.1101/505958

**Authors:** Simona Olmi, Spase Petkoski, Maxime Guye, Fabrice Bartolomei, Viktor Jirsa

**Affiliations:** Inria Sophia Antipolis Méditerranée Research Centre, MathNeuro Team, 2004 route des Lucioles-Boîte Postale 93 06902 Sophia Antipolis, Cedex, France; CNR – Consiglio Nazionale delle Ricerche – Istituto dei Sistemi Complessi, 50019, Sesto Fiorentino, Italy; Aix Marseille Université, Inserm, Institut de Neurosciences des Systèmes, UMR_S 1106, 13005, Marseille, France; Faculté de Médecine de la Timone, centre de Resonance Magnétique et Biologique et Medicale (CRMBM, UMR CNRS-AMU 7339), Medical School of Marseille, Aix-Marseille Universite, 13005, Marseille, France; Assistance Publique – Hôpitaux de Marseille, Hôpital de la Timone, Pole d’Imagerie, CHU, 13005, Marseille, France; Assistance Publique – Hopitaux de Marseille, Hopital de la Timone, Service de Neurophysiologie Clinique, CHU, 13005 Marseille, France

## Abstract

Information transmission in the human brain is a fundamentally dynamic network process. In partial epilepsy, this process is perturbed and highly synchronous seizures originate in a local network, the so-called epileptogenic zone (EZ), before recruiting other close or distant brain regions. We studied patient-specific brain network models of 15 drug-resistant epilepsy patients with implanted stereotactic electroencephalography (SEEG) electrodes. Each personalized brain model was derived from structural data of magnetic resonance imaging (MRI) and diffusion tensor weighted imaging (DTI), comprising 88 nodes equipped with region specific neural mass models capable of demonstrating a range of epileptiform discharges. Each patient’s virtual brain was further personalized through the integration of the clinically hypothesized EZ. Subsequent simulations and connectivity modulations were performed and uncovered a finite repertoire of seizure propagation patterns. Across patients, we found that (i) patient-specific network connectivity is predictive for the subsequent seizure propagation pattern; (ii) seizure propagation is characterized by a systematic sequence of brain states; (iii) propagation can be controlled by an optimal intervention on the connectivity matrix; (iv) the degree of invasiveness can be significantly reduced via the proposed seizure control as compared to traditional resective surgery. To stop seizures, neurosurgeons typically resect the EZ completely. We showed that stability analysis of the network dynamics, employing structural and dynamical information, estimates reliably the spatiotemporal properties of seizure propagation. This suggests novel less invasive paradigms of surgical interventions to treat and manage partial epilepsy.

PACS numbers:

**AUTHOR SUMMARY:** Epilepsy is characterized by perturbed dynamics that originate in a local network before spreading to other brain regions. We studied patient-specific brain network models of epilepsy patients, comprising 88 nodes equipped with region specific neural mass models capable of demonstrating epileptiform discharges. Applying stability analysis led to a seizure control strategy that is significantly less invasive than the traditional surgery, which typically resects the epileptogenic regions. The invasiveness of the procedure correlates with graph theoretical importance of the nodes. The novel method subsequently removes the most unstable links, a procedure possible by advent of novel surgery techniques. Our approach is entirely based on structural data, allowing creation of a brain model based on purely non-invasive data prior to any surgery.

## II. INTRODUCTION

Propagation of activity through a network is a non-stationary spatiotemporal process and is the most fundamental representation of information processing in the brain [1, 2]. In task conditions, as the behavioral dynamics unfold, brain activity simultaneously evolves in a hierarchy of characteristic network activations. For instance, decision making tasks involve a chain of sequential subnetwork activations comprising the initial preparatory phase, the decision phase and the final decision execution phase [3, 4]. Information processing in sensorimotor coordination [5] and auditory, visual and linguistic tasks [6] show robust propagation through well-timed activation chains of characteristic subnetworks.

In the diseased and aging brain, the spatiotemporal organization of information processing is disrupted [7, 8]. Local and distributed alterations of connectivity and cell tissue properties influence the functional capacity of the brain in stroke, schizophrenia, multiple sclerosis and a range of neurodegenerative disorders [9–13]. From the perspective of spatiotemporal signal propagation and cognitive networks, epilepsy takes a key role, since seizure propagation is often accompanied by the evolution of a characteristic behavioral pattern, the semiology, which unfolds behaviorally as the neuroelectric activity is traced out in space and time. Epilepsy is a common disorder, affecting over 65 million people worldwide [14], frequently accompanied by cognitive and memory deficits [15, 16]. The investigation of electrographic spatiotemporal patterns (including interictal spikes, rhythmic bursts, wave propagation) relative to cognition networks dates back to 1939 [17] and is a crucial step in improving treatment and quality of life of patients with epilepsy.

The influence of network topology and the anatomical organization of the epileptogenic process are particularly important in the context of seizure control and epilepsy surgery [18]. Bancaud and Talairach called the region of seizure onset the “epileptogenic zone” (EZ) [19], which is not necessarily identical to the structural lesion and seizures have been reported to arise from structures distant from the lesion itself [19]. Often the clinical manifestations do not occur with seizure onset, but actually with the propagation of the seizure to other regions, which places further importance upon the understanding of how the spatiotemporal organization and spread of discharge activity arises. The concept that focal epilepsies are indeed not so local and involve large scale macroscale circuits is becoming increasingly more accepted in epileptology [18, 20–24].

Practically, however, there are fundamental obstacles to the application of network control in the spatiotemporal organization and propagation of healthy and pathological activity. Antiepileptic drugs are most common and act globally by mostly targeting ion channels and inhibitory neurotransmission and synaptic receptors across the entire brain network. Yet approximately 30% of epilepsy patients suffer from drug-resistant epilepsy and continue to experience seizures despite appropriate antiepileptic drug treatment [25]. For patients with medically refractory epilepsy there are two possible control techniques: resective surgery and neurostimulation, where for the latter the epileptogenic zone is electrically stimulated in order to suppress seizures. In recent years, cortical and subcortical neurostimulation is emerging as a promising treatment for drug-resistant epilepsy [26, 27]. On the other hand, regarding the surgical removal of brain tissue, the most common network control technique consists in removing the epileptogenic zone (ascribed to temporal and extratemporal resections and hemispherectomy). The second, less common type of epilepsy surgery interrupts nerve pathways that allow seizures to propagate. This procedure, termed disconnection, is used for corpus callosotomy, hemispherotomy and multiple subpial transections [28]. The application of disconnection to other brain regions has been largely unsuccessful [28] and the causes of failure remain unknown. With the advent of novel minimally invasive surgery techniques, microsurgical disconnections become realistic to be performed in the human and present the least invasive restorative intervention for brain network control [29]. These tech niques include laser surgery via thermal ablation and Gamma knife radio surgery, in which converging narrow ionizing beams are focused to destroy cell tissue. Whether the goal of the network intervention is the suppression of seizure activity or alleviation of cognitive deficits due to comorbidities, conditio sine qua non for this will always be a good understanding of the consequences of the network manipulation.

Successful outcomes of either treatment depend critically on the ability to accurately identify the epileptogenic zone, which is a long, complex, and costly process. In the case of intracranial electrocorticographic (ECoG) recordings the skull has to be opened through a craniotomy, thus allowing the surgical implantation of electrodes into the patient’s brain. Afterward the patient undergoes prolonged and costly monitoring, with the risk of contracting infection and neurological comorbidities, while the clinical team waits for several seizure events to accrue sufficient EEG data. Manual inspection of the signals finally allows specialized epileptologists and technicians to localize the epileptogenic zone. However surgical outcomes are highly variable irrespectively of the presurgical evaluation via invasive recordings: between 30%-70% of patients continue having seizures 6 months after treatment, which may be due to mislocalization of the epileptogenic zone. To overcome this problem, scientists have done an intensive search for an accurate data analytics tool to reduce time, risks and costs of invasive monitoring. To address this issue in models and patient brains, we here construct patient-specific connectome-based large-scale brain network models and identify the least invasive strategies to stop seizure propagation. Personalized connectomes are composed by the structural links derived from diffusion weighted tensor imaging (DTI) and tractography, and have been shown to predict individual functional connectivity better than generic approaches. Empirical studies are impossible to be performed systematically in the human and are limited to sparse sampling (empirical realizations) only. In silico modeling allows to complete the analysis with high-density sampling (all possible in-silico realizations) and identify the best network modulation for network control. We adapt a disconnection approach, aiming to identify the minimal ablation for a particular patient to stop the propagation of activity through the personalized network, let this be propagation due to stimulation or spontaneous seizures. Neuroelectric discharges are captured by gold standard data of epileptic patients recorded via intracranial electrodes. Brain areal dynamics is derived from the mean activity of neural populations via neural mass models able to capture the details of the autonomous slow evolution of interictal and ictal phases. Personalized brain network models are then derived for 15 patients. We systematically investigate all possible realizable disconnections of the epileptogenic zone and demonstrate that for all patients a partial targeted disconnection, accounting for the resection of one or a few pathways, is sufficient to stop seizure propagation in the brain, systematically reducing invasiveness and outperforming generic (non-personalized) procedures. In particular partial targeted disconnection is designed by combining the analysis of patient specific structure with model dynamics and performing a Linear Stability Analysis of the system. The preferred pathways along which the (infinitesimal) perturbations spread, turn out to be fundamental to identify the propagation and recruitment process of a seizure, thus suggesting the virtual ablation to perform in the personalized connectomes.

## III. RESULTS

During an epileptic seizure the brain shows an abrupt onset of fast oscillatory activity, which typically propagates to other regions and recruits these as oscillatory subnetworks. The seizure originates in the Epileptogenic Zone (EZ) and propagates to the Propagation Zone (PZ) (see Fig. 1). The recruitment of the regions in the propagation zone can happen either by independent activation of the single areas (Fig. 1, time *t*_2_), or by activating multiple areas at the same time (Fig. 1, times *t*_3_ – *t*_5_). The distinction of EZ and PZ based on the electrographic signatures is non-trivial, in particular, when the spreading continues after the epileptogenic zone has seized its intense activity (Fig. 1, time *t*_6_). In case of asymptomatic seizures, the activity is often limited to the epileptogenic zone and no propagation takes place.

**Figure 1:**
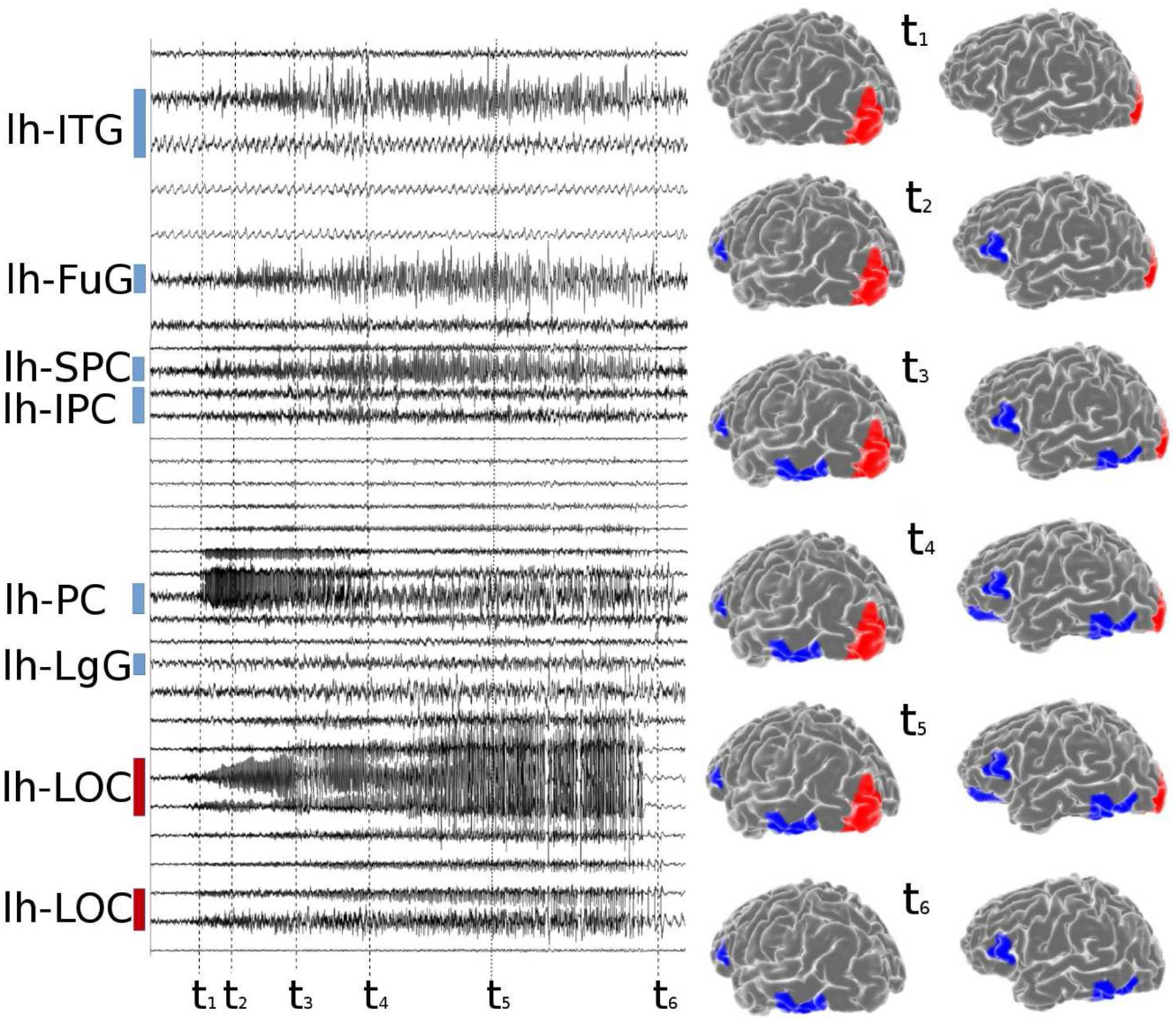
Stereotactic Electroencephalographic (SEEG) Data of patient CJ. Left panel: time series of partial seizures recorded with SEEG. The colored bars indicate the EZ (red)and PZ (blue). Right panel: On the top, the spatial organization of the EZ and PZ is shown on the left hemisphere, frontal-lateral and lateral view. Below, spatiotemporal activation patterns are plotted at different time points of the seizures. The same time points highlighted with black dashed lines in the left panel. lh-SPC, left superior parietal cortex; lh-IPC, left inferior parietal cortex; lh-LgG, left lingual gyrus; lh-LOC, left lateral occipital cortex; lh-FuG, left fusiform gyrus; lh-PC, left Pericalcarine; lh-ITG, left inferior temporal gyrus.

### A. Comparison among different methodologies

Once the EZ is schematized as one (or more) nodes inside the brain network, then the most common epilepsy surgery actually performed, which is ascribed to the resection of the epileptogenic zone, corresponds to the disconnection of all the pathways outgoing the focal nodes in addition to the removal of the node itself. Fig. 2 shows the connectivity matrix of an epileptogenic brain (light blue links) with the EZ consisting of one area, where the set of connections outgoing the focus is highlighted in blue and pink. The targeted disconnection method aims at interrupting only those nerve pathways that allow a seizure to propagate and corresponds in this framework to the resections of the few pink links outgoing the EZ selected via the Linear Stability Analysis procedure. In Fig. 3 we compare the removal of: i) all the links outgoing the epileptogenic zone (corresponding to the standard clinical resection); ii) a set of randomly chosen links outgoing the epileptogenic zone; iii) the strongest links outgoing the macro-area composed by epileptogenic and propagation zones; iv) the strongest links outgoing the epileptogenic zone; v) links selected via the Linear Stability Analysis procedure. Here strongest links refer to the connections with largest weights. Since the connectivity matrix is normalized so that the maximum value is one, the strongest links are of order O(1), while are not considered as strongest links those with weights of order of magnitudes o(1). From a computational point of view the virtual ablation of a link corresponds, in all the mentioned cases, to the removal of the corresponding row and column in the structural connectivity matrix of the investigated epileptogenic brain.

**Figure 2:**
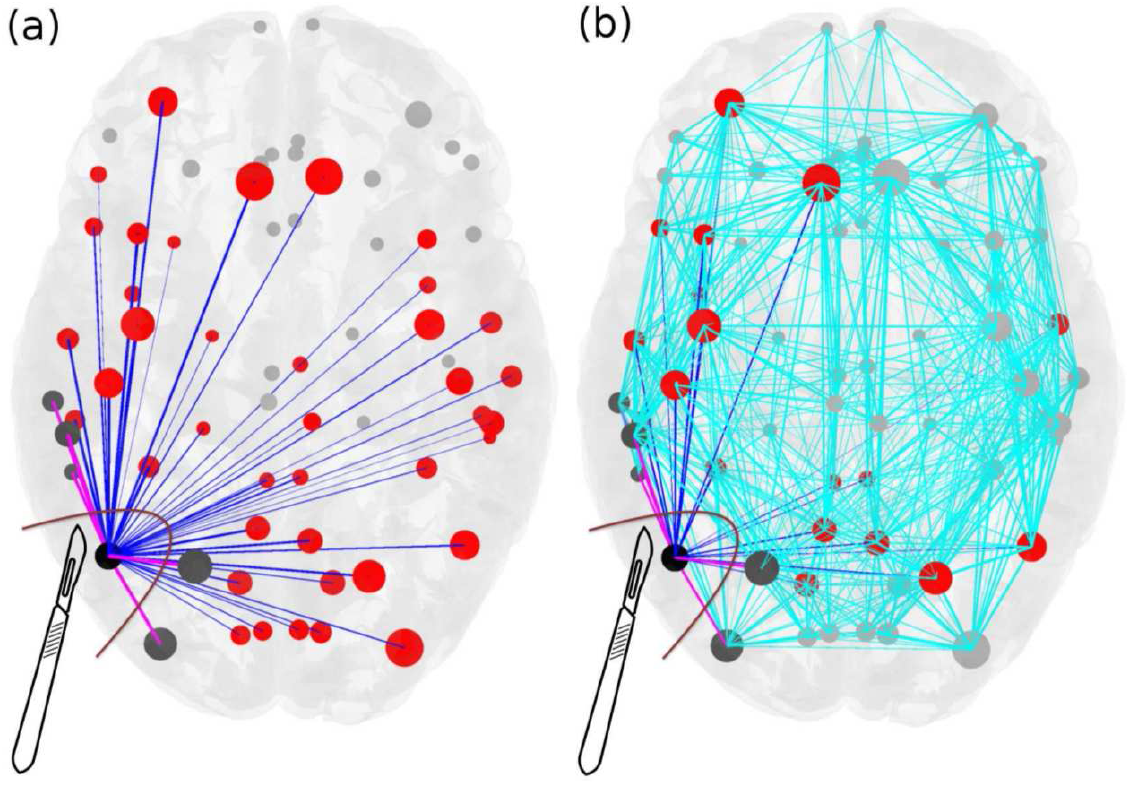
Scheme for comparison of standard resection methods, where the entire epileptogenic zone (EZ) is removed during surgical operation, versus lesioning minimal number of links. Panel (a): The connectivity matrix is illustrated for an epileptogenic brain with the EZ consisting of one area (black). The outgoing connections of the EZ are blue and pink, connecting red and dark grey areas respectively, and they are all removed during the current surgical procedures of disconnecting the EZ. Targeted lesioning depicts the minimal number of links that are sufficient to be removed (pink) in order to stop the seizure, versus the total number of outgoing links from the EZs (blue) that are removed during the resection of an entire EZ. Light blue links added in panel (b) represent the full connectivity of the network. The size of the nodes reflects how strongly they are connected, and the width of the links correspond to their weight. Unaffected nodes by any of the resection procedures are light grey and unaffected links are light blue.

**Figure 3:**
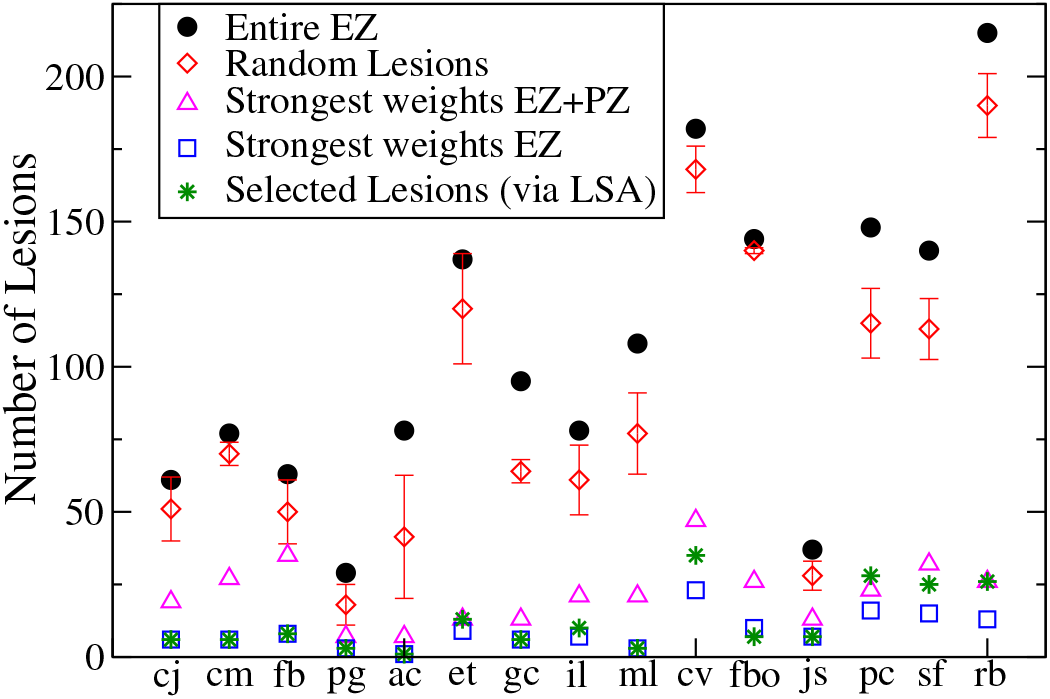
Comparison between the results obtained with 5 different lesioning procedures for each of the 15 analyzed patients. In particular it is shown the numbers of links that are cut in order to stop the seizure propagation, if i) the entire EZ is removed (black dots); ii) random cuts are done (red diamonds); iii) the strongest links outgoing the macroarea (EZ + PZ) are removed; iv) the strongest links outgoing the EZ are cut; v) selected lesions are done following Linear Stability Analysis indications (green stars). The data for the case (ii) are calculated averaging over 5 different realizations of random lesioning procedures. Patients are ordered according to the extension of their EZ: the focus is represented as a single node for patients cj-pg; 2 nodes for patients ac-ml; 3 nodes for patients cv and fbo; 4 nodes for patients jc-sf; 6 nodes for rb.

The random choice of the links to be cut among the subgraph connecting the EZ with the rest of the network (case ii), represents the mathematical procedure that better approximate the results of standard clinical resections, where all the subgraph is removed, since a big amount of connections needs to be resected in order to stop the seizure propagation and to prevent the emergence of epileptic symptomatic seizures. On the other hand, if we restrict our attention to the connections outgoing the region composed of both EZ and PZ (case iii), the strength of the links in this macro-area might be *a priori* a good indicator to understand the role played by the topology in the spreading of the seizure all along the brain. Actually, the mechanism underlying the seizure propagation is more subtle and the topological features of the structural connectivity matrices are not sufficient themselves to fully explain the recruitment of other areas occurring in the propagation. This means that if we try to stop the seizure propagation cutting the strongest links outgoing the macroarea EZ+PZ, the number of fundamental links that we identify according to this procedure are more than the ones actually responsible for the mechanism. When we apply the same criterion to the subgraph outgoing the EZ only (case iv), thus disconnecting the strongest links, we are able indeed to design an effective targeted disconnection. Due to the distribution of the weights of the structural connectivity matrices, connections with biggest weights turn out to be easy to recruit. The least invasive method identifying the minimal number of pathways, along which the seizure propagates, is the Linear Stability Analysis (case v), that univocally identifies, via the calculation of the stability of the system in presence of perturbations (represented by the seizures), the most unstable directions along which the recruitment and the seizure spreading take place. These most unstable directions, that are represented by the links connecting different populations, are often the strongest links outgoing the epileptogenic zone. In particular this is always true when the epileptogenic zone is composed by a single area, i.e. a single node.

However, it is worth noticing that purely structural information is not sufficient to predict the propagation and eventual stopping of the seizure, and the availment of a mathematical model is required to predict correctly the least invasive intervention. To demonstrate this, we compute for the structural connectivity matrix of a specific patient (CJ), various network measures, including efficiency, strength, betweeness, centrality and clustering (see Figs. S2-S4, Text S2), which show that neither the epileptogenic zone nor the propagation zone are characterized by outstanding values of these indicators. In general, epileptogenic and propagation zones have intermediate values of all the graph metrics. Thus, if we concentrate on the information gained by the network theory only, analyzing the structural connectivity matrix via the identification of the strongest links and the calculation of the network measures, it would be impossible to predict the number of recruited populations in the seizure spreading process. This is because not only the identification of the propagation zone but also the choice of the strongest links in the connectivity matrix are not well-defined due to the fact that strong links are present even outside the EZ. Therefore, as a first step, a dynamical model that describes the activity of each node (see eqs. 1) is necessary to verify the self-emergent behavior in the network and to test the procedures i)-iv) shown in Fig. 3. However, a general procedure, independent on the specific connectivity matrix, is needed to investigate the response of the system under perturbations and to identify the propagation direction of the instabilities related to the seizure propagation. Actually the Linear Stability Analysis offers a way to calculate the stability of the system against perturbations and to identify the directions along which the system is less stable combining the information on the connectivity matrix with the information on the dynamics (see eqs. 2). Thus, the targeted lesion procedure, identified with the Linear Stability Analysis, is to be preferred to the analysis of the strongest links only, since it is able to give insights on the dynamics of the system and its criticalities, even though its predictions are comparable with those given by the procedure iv).

### B. Detailed analysis of a single patient

From the computational point of view, the dynamics of each node of the network parcellation is given by a neural mass model able to reproduce the temporal seizure dynamics and to switch autonomously between interictal and ictal states [30]. Nodes are connected together via a permittivity coupling acting on a slow time-scale, that is on the time-scale of seconds, which is sufficient to describe the recruitment of other brain regions [31] (see Methods and Texts S1, S2). The results, presented hereafter, are obtained for the structural connectivity matrix of patient CJ, a patient with Engel score [32] III (see Text S3 and Figs. S5-S10 for the results related to other patients, with Engel scores I-III). In particular the self-emergent dynamics of this system is shown in Fig. 4 (upper panels). The EZ, chosen according to the medical doctors’ evaluation, is the Lateral occipital cortex (LOC), placed on the left hemisphere. The seizure onset is followed by the recruitment and successive seizure emissions of the PZ: Fusiform gyrus (FuG), Superior parietal cortex (SPC), Inferior temporal gyrus (ITG), Inferior parietal cortex (IPC), Pericalcarine (PC), Lingual gyrus (LgG), all located in the left hemisphere (upper left panel). The PZ, which represents the dominating sub-networks involved in the transition towards the seizure state, is determined self-consistently once the EZ is given (see Methods and Text S1) and differs partly from the one predicted by the clinicians on the basis of EEG and SEEG signals. In particular the clinical PZ prediction identified Inferior parietal cortex and Superior parietal cortex as dominant subnetwork, while Fusiform gyrus and Inferior temporal gyrus were not taken into account during the presurgical evaluation. Moreover, once the pathways to be resected are chosen via the Linear Stability Analysis suggestions and the targeted disconnection is performed, the recruitment process is no longer possible in the system even though the EZ is still triggering seizures (Fig. 4 upper right panel).

**Figure 4:**
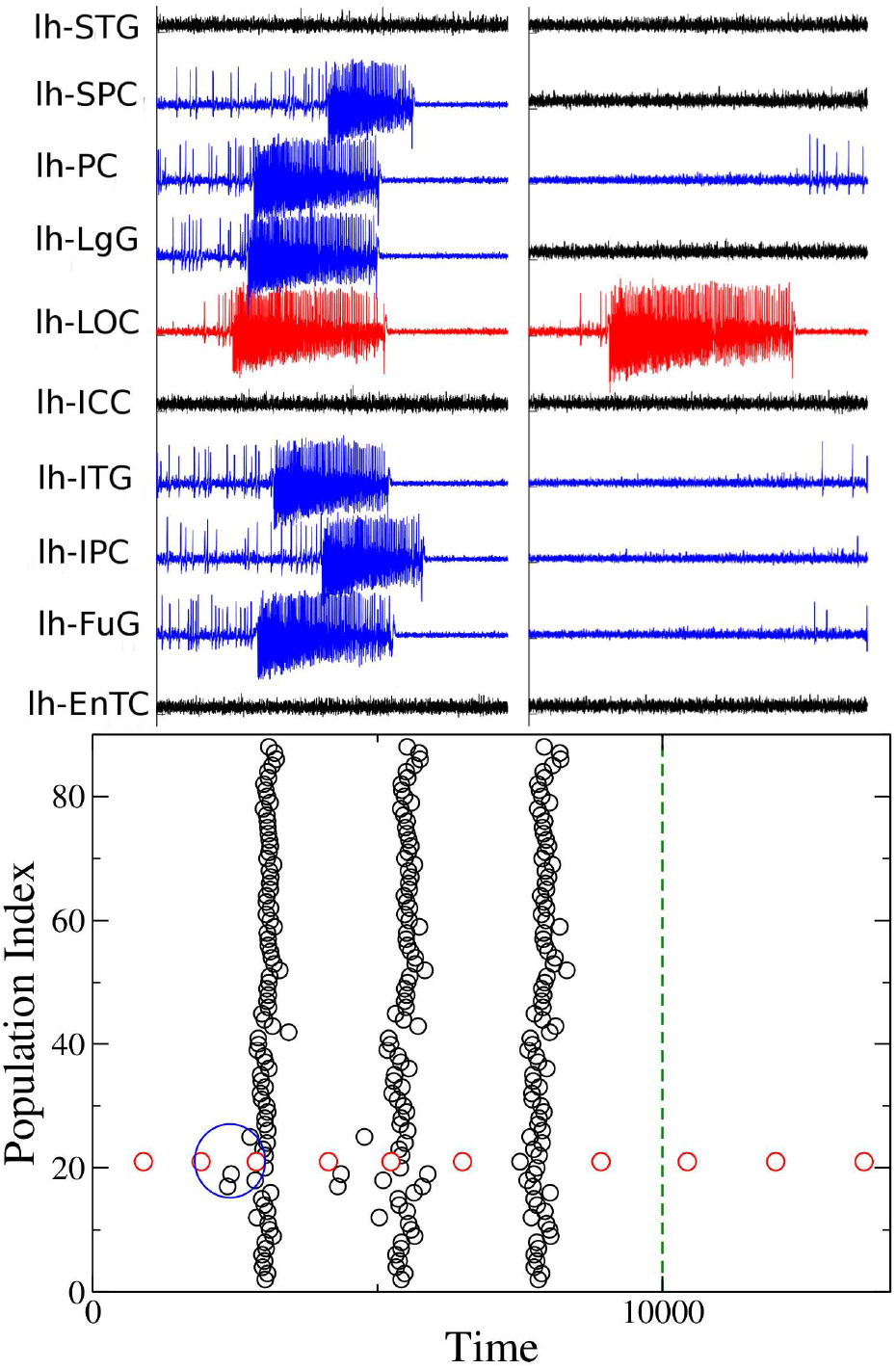
Analysis of patient CJ. Clinical history: occipital epilepsy type (left size). EZ region: Lateral occipital cortex. PZ prediction: Fusiform gyrus, Superior parietal cortex, Inferior temporal gyrus, Inferior parietal cortex, Pericalcarine, Lingual gyrus. PZ clinical prediction: Inferior parietal cortex, Superior parietal cortex. Lesions: links between regions LOC-FuG, LOC-SPC, LOC-ITG, LOC-IPC, LOC-PC, LOC-LgG must be cut in order to stop the seizure (lesions are performed in correspondence of the time identified by the dashed blue line). Upper panels: Time series generated by the implemented brain network model with the connectome of patient CJ. On the left the PZ (blue curves) is recruited immediately after the seizure (red curve) is emitted; on the right the recruitment is no more possible after the targeted disconnection is performed. Lower panel: Seizure events as a function of time. The connectivity matrix consists of 88 nodes and the EZ corresponds to 21. Seizures emitted by node 21 are highlighted in red, while the others are given in black. Green dashed line corresponds to the time at which lesions occur. The blue ellipse highlights the moment in time at which the PZ is recruited after the seizure emitted by the EZ.

A different visualization of the seizures onsets, taking place in the network at different times, is shown in Fig. 4 (lower panel): each seizure emitted by a single node (EZ) at a specific time is identified with a black (red) dot, as in a spike raster plot, which is a typical representation of neural spike-timing activity. In this way it is possible to have an overview of the entire network for long time periods. In particular the epileptogenic zone is represented in the raster plot as node 21 and its emitted seizures are identified with red dots for consistency with the upper panels. EZ is able to trigger seizure autonomously and, after a small time interval, to recruit a small number of regions: the recruitment event highlighted in the blue circle is the same as shown in the upper left panel. After a small delay, suddenly, the seizure is no more confined to a small number of regions, but it is able to propagate along the entire network, and all regions start emitting seizures synchronously (as in a bursting activity). This mechanism is self-sustained unless selected lesions are performed. In particular, at the time corresponding to the blue dashed line in Fig. 4 (lower panel), six lesions are performed on the structural connectivity and the seizure is no longer able to recruit any sub-network. The ongoing dynamics after the disconnection is performed, is the same shown in Fig. 4 (upper right panel). The lesions correspond to cutting the connections between the EZ and the nodes in the PZ that are involved in the initial recruitment phase and are identified by calculating the maximal eigenvector of the Jacobian matrix (see Methods and Text S1).

The dynamics presented in Fig. 4, can be also characterized in terms of the eigenvalues shown in Fig. 5 (a). In particular, the dynamics describing the seizures of patient CJ, is characterized by 2 positive eigenvalues, thus meaning that the system in unstable. The eigenspectrum changes if some topological modifications (e.g. lesions) are implemented into the original structural matrix. In particular the number of positive eigenvalues diminishes by each further removal of the links, following the Linear Stability Analysis procedure. It is important to notice that the necessary links to be cut in order to stop the seizure are those between the EZ (node 21) and the first six most unstable nodes. Once these links are removed the system is still unstable, because the epileptogenic zone is still active, but the epileptogenic activity of this area is no more able to recruit other zones.

**Figure 5:**
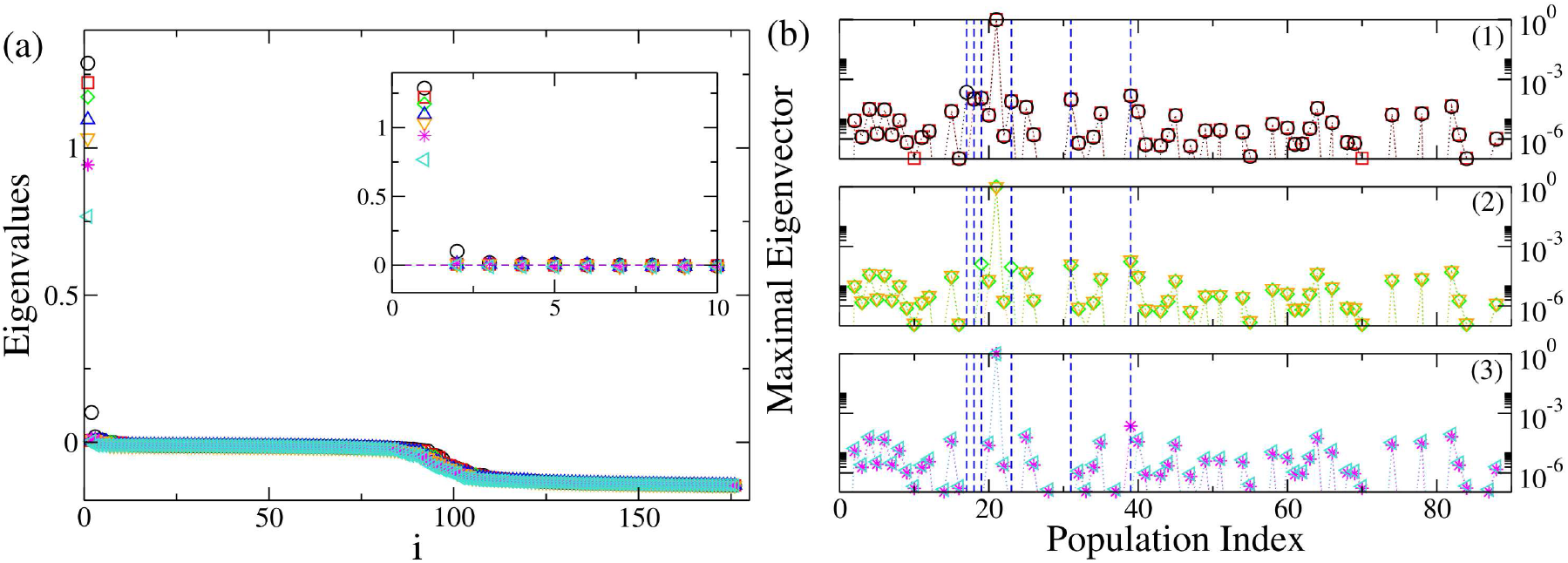
Analysis of patient CJ. Panel (a): Eigenvalues of the system when the original structural connectivity is considered as a network structure (black circles). The index i on the x-axis represents the eigenvalues index (for more details see SI). Red squares represent the eigenvalues of the networks when the link between the populations (nodes) LOC-FuG (21-17) is removed. Green diamonds represent the eigenvalues of the networks when the links between the populations (nodes) LOC-FuG, LOC-SPC (21-17, 21-39) are removed. Blue triangles represent the eigenvalues of the networks when 3 connections between the regions (nodes) LOC-FuG, LOC-SPC, LOC-ITG (21-17, 21-39, 21-19) are removed. Orange triangles represent the activity when 4 connections between the regions (nodes) LOC-FuG, LOC-SPC, LOC-ITG, LOC-IPC (21-17, 21-39, 21-19, 21-18) are removed. Magenta stars: links between the regions (nodes) LOC-FuG, LOC-SPC, LOC-ITG, LOC-IPC, LOC-PC (21-17, 21-39, 21-19, 21-18, 21-31) are removed. Turquoise triangles: links between the populations (nodes) LOC-FuG, LOC-SPC, LOC-ITG, LOC-IPC, LOC-PC, LOC-LgG (21-17, 21-39, 21-19, 21-18, 21-31, 21-23) are removed. In the inset is shown an enlargement of the first, positive part of the spectra. Panel (b): Maximal eigenvectors calculated when the original structural matrix is considered as network structure and when successive links are removed. Symbols and the color code are the same as in panel (a). The blue lines represent the most unstable localized nodes to which are associated the most unstable directions. In particular panel (b1) shows the situation where the structural matrix is intact (black circles) and where the first link has been removed (red squares). Removing the link between the populations (nodes) LOC-FuG (21-17) corresponds to the loss of importance of node 17, which is not localized anymore. In panel (b2) are shown the maximal eigenvectors in case 2 removals (green diamonds), 3 removals (blue triangles) and 4 removals (orange triangles) are operated. Every time an additional link is cut according to the Linear Stability Analysis procedure detailed for panel (a), the corresponding node does not appear any more as important (i. e. localized) when calculating the maximal eigenvector. Finally in panel (b3) are shown the maximal eigenvectors in case 5 removals (magenta stars) and 6 removals (turquoise triangles) are operated: when all the relevant lesions are performed there are no more localized nodes (except the EZ), thus suggesting that the unstable directions are not accessible anymore.

In Fig. 5 (b), the procedure for identifying the most unstable nodes is described in more details. Firstly, we show the maximal eigenvector calculated for different network configurations. The maximal eigenvector is associated with the maximum positive eigenvalue resulting from the calculation of the Jacobian matrix. It is important to notice that in this calculation is taken into account the connectivity of the specific patient. Therefore, when the original structural connectivity matrix is considered as network structure, the largest elements of the eigenvector correspond to some populations in the brain parcellation and the most unstable directions turn out to be the connections between these populations and the EZ. The largest elements of the eigenvector are indicated by blue dashed lines in Fig. 5 (b); the biggest one (non indicated with a dashed line) is the EZ. In particular, if the link between the epileptogenic population and the second largest element (corresponding to FuG – node 17) is cut (sub-panel (1), red squares), this direction is no longer unstable and the node looses its importance, as element of the maximal eigenvector, no longer having a relevant amplitude. In particular the entry corresponding to the node, whose connection has been resected, is now characterized by a very small absolute value, while other directions may be enhanced. The other largest elements of the vector (indicated by blue lines) still constitute the set of most unstable directions, while this is no longer true for the resected connection LOC-FuG (21-17), related to the entry number 17 of the maximum eigenvector. This discussion applies to the first 6 biggest elements of the eigenvector. Once all the 6 links are cut (see sub-panel (3), turquoise triangles), the eigenvector is no more localized in specific zones. The EZ is the only one left significantly larger than zero, but it is no longer contributing to the instability of the system in a relevant way.

### C. Systematic analysis

The former analysis has been done on the basis of the medical doctors’ prediction for the EZ. However it is worth doing a systematic analysis of the network dynamics, by supposing that the EZ is placed in every possible node of the connectome. In particular, by taking into account the structural connectivity matrix of patient CJ and by systematically placing the EZ in different nodes, we investigate the dynamics of the full network for all possible epileptogenic nodes. For every shuffled position of the EZ, the Linear Stability Analysis is calculated, henceforth the eigenvalues and eigenvectors of the system are obtained for all these configurations. Even though the results depend on the position of the EZ, it is possible to demonstrate that, in general, by performing a virtual resection of a small number of pathways, up to a maximum of 15, the seizure propagation can be stopped for any initial EZ considered (see Fig. 6(a)). This proves that the performed lesioning procedure is independent on the chosen particular case and it can be validated for general EZ locations.

The lesioning procedure for a given structural connectivity network directly affects the size of the PZ. As expected, it shrinks when increasing the number of removed links. This is a natural consequence of the fact that the number of possible unstable directions decreases for every resection, and the number of regions that are likely to be recruited vanishes. More specifically, the PZ generally shrinks to 0-2 areas, after 5-6 performed disconnections (see Fig. 6(b)). In most of the cases this is sufficient to alter the seizure spreading, since a small PZ, made of 2 populations maximum, is not able to convey the spreading: the most unstable directions are no more accessible by the system. In this case the system undergoes a so-called asymptomatic seizure.

**Figure 6:**
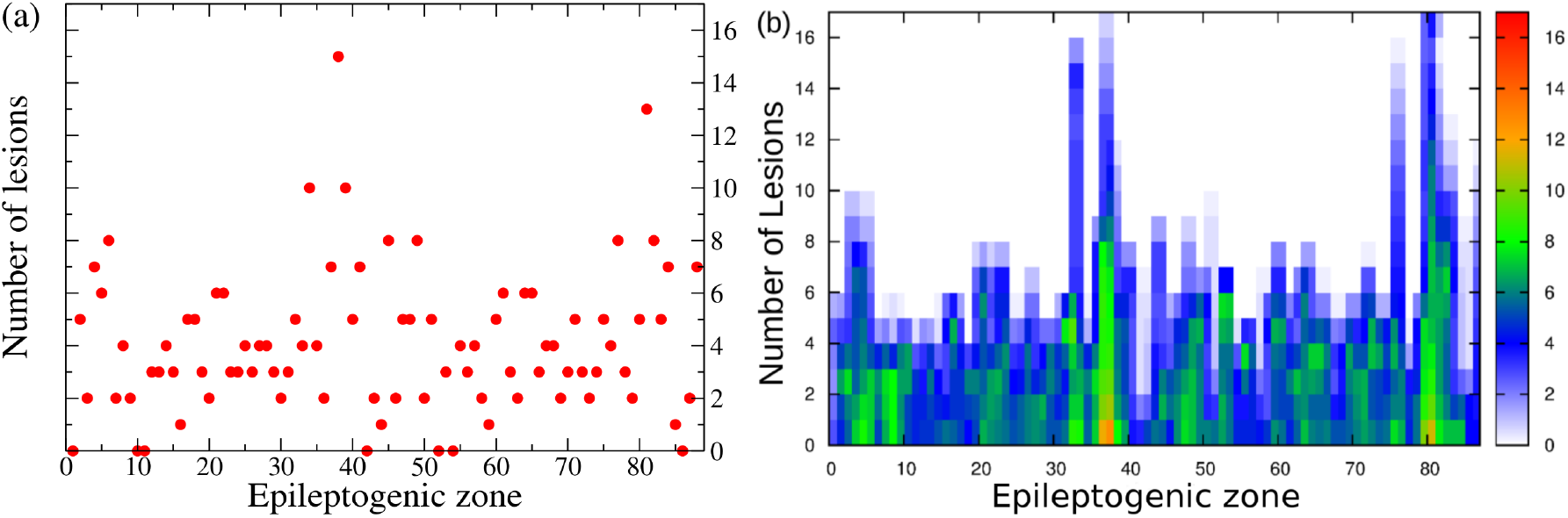
Panel (a): Number of lesions necessary to stop seizure propagation as a function of the epileptogenic region. Panel (b): Dependence of the PZ size on each lesion, as a function of the EZ. The color code indicates the number of regions belonging to the PZ, from 0 (white pixel) to 17 (red pixel).

A topological analysis *a posteriori* on the dependence of the partial targeted disconnection on the different graph connectivity measures, highlights the significance of these quantities. Namely, starting from the results reported in Fig. 6(a), we have further calculated the dependence of the disconnection procedure on efficiency, strength, clustering, degree, betweenness, centrality (see Fig. 7). In particular these topological metrics have different values for different choices of the EZ. The results indicate that the number of needed topological interventions on the network are strongly proportional to the efficiency, clustering and betweenness, whereas, while still present, this relationship is less pronounced for the strength, degree and centrality. This means that controlling the seizure spreading is more difficult when the involved EZ belongs to a dense, high-clustered neighborhood. In this situation each node involved can easily transmit information among the others: a clique of regions that rapidly communicate by using a dense subgraph plays a big role in controlling the network and enhancing the seizure propagation. Once the clique controls the dynamics, it is necessary to cut more links responsible for the information flow to propagate. On the contrary, the shortage of effective communication networks among the pathological areas facilitates the control of the network, and in this case the goal can be achieved with a much smaller number of topological interventions.

**Figure 7:**
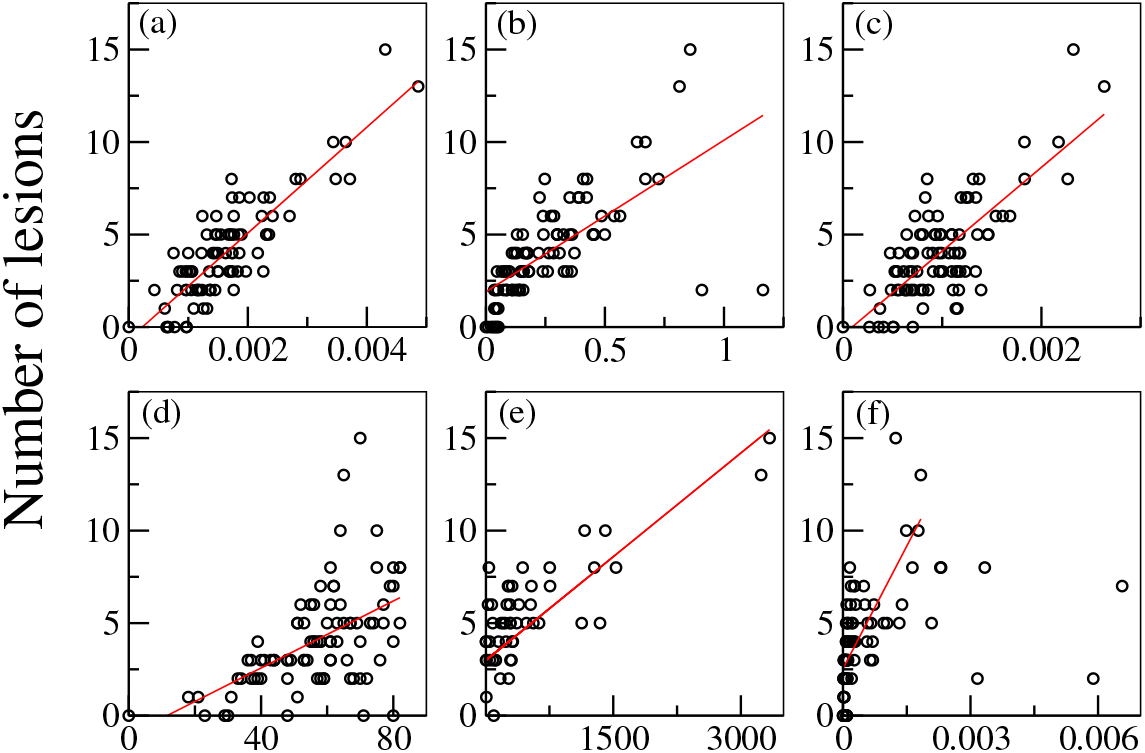
Number of lesions needed to stop the propagation for different graph measures of each EZ, calculated for the connectivity matrix of the patient CJ. Graph metrics reported on the x-axis are: (a) efficiency; (b) strength; (c) clustering; (d) degree; (e) betweenness; (f) centrality. The red line in each panel shows the correlation among the data, and the estimated Pearson correlation coefficient is: (a) 0.8615377; (b) 0.6735522; (c) 0.765353; (d) 0.540695; (e) 0.7852488; (f) 0.7165724. All p values of statistical significance are below 10^−6^.

The importance of separately considering the structural connectivity matrices of single patients, for better designing the partial targeted disconnection, can be illustrated comparing the previous results with the ones obtained by performing the Linear Stability Analysis procedure on an average connectome. The latter is constructed by averaging the 15 structural connectivity matrices of the considered patients (via an algebraic mean). It turns out that the analysis on the average connectome is much less predictive and that the individual variability has clear effects upon the outcome of the prediction results, and therefore, of possible therapies and treatment approaches. In particular, the resulting average connectome has different topological characteristics with respect to the matrices of the single patients: the degree of each node is larger and in general, it is more dense and homogeneous. If we select as EZ, the area suggested by the clinical prediction (i. e. node 21, lh-LOC for the patient CJ) and we calculate the maximum eigenvector, it turns out to be no longer localized, like the one shown in Fig. 6 (b). Now the eigenvector has a distributed structure, with many elements having a considerably large absolute value (i. e. above 10^−5^). Hence, it is more difficult to individuate a small number of nodes that actively participate to the dynamics and that are overwhelmingly involved in the seizure propagation. On the contrary, many more nodes are now involved in the propagation, and many more directions are viable for the spreading of the pathological activity. A comparative analysis of the number of links that must be cut for stopping the seizure propagation, obtained by following different procedures, is reported in Table 1.

**Table I:** Comparison of the minimum necessary number of lesions for stopping the seizure propagation for different lesioning procedures.

| Type of analysis | Lesions |
| --- | --- |
| Linear Stability Analysis (LSA) on the Patient's Structural Connectivity Matrix (PSCM) | 6 |
| Strongest Links in the PZ (on the PSCM) | 22 |
| Random Cuts (on the PSCM) | 51 |
| All the outgoing links of the EZ (on the PSCM) | 61 |
| LSA on the Average Connectome | 66 |
| Random Cuts on the Average Connectome | 69 |
| All the outgoing links of the EZ on the Average Connectome | 71 |

While it is sufficient to cut only 6 links detected via the Linear Stability Analysis procedure on single patient’s connectome, it is necessary to remove up to 66 links before the seizure stops propagating on the averaged connectome, even thought the same procedure have been used. The latter number is comparable with the results obtained by random removal of the links belonging to the subgraph of the EZ in the patient’s connectome, and hence it is no better than a “blind” resection.

## IV. DISCUSSION

Connectivity is fundamental to the transmission of information through a network, let it be pathological or physiological. With regard to the mechanisms underlying propagation in networks, several mechanisms have been hypothesized. A basic form of propagation is energy dissipation [33], which depends on the physical properties of the network such as the local response of the tissue and its connectivity. Following a local stimulation, the brain network shows initially the well-known local propagation via the intracortical connections within the grey matter, but then the spatiotemporal organization after 300 msec is dominated by the large-scale white matter connection topology [33]. As this process is independent of any representation of function, propagation within cognitive networks rely at least implicitly on some notion of coding, that renders an activation profile functionally meaningful. For instance, increased interareal gamma-band coherence and phase synchronization have been hypothesized as markers of selective attention allowing different input information streams to be routed through brain networks according to task relevance [2, 34, 35]. In the diseased brain the propagation of this information is disturbed. Seizure propagation, in particular, is generally based on the concept of abnormalities of synchronizability: i) seizure evolution is driven by the strong synchronizing activity of the EZ which is able to guide the surrounding tissue [36]; ii) the surrounding tissue has a diminished ability to contain and regulate abnormal activity, and it allows the seizure to propagate [37]; iii) synchronizing and desynchronizing nodes operate antagonistically, such that synchronizability increases and dynamical processes diffuse easily through the network when synchronizing nodes exert greater push than desynchronizing nodes [38].

In all these cases of propagation, the frame, in which the activity evolves, is defined by the energy dissipation properties of the network and can be controlled and rerouted through network rewiring, of which the disconnection type is the simplest to realize, even though in clinical practice it has not found many applications so far. The few existing applications exclusively rely on a complete disconnection such as corpus callosotomy or hemispherotomy, since in absence of understanding the network effects upon propagation, more subtle and minimally invasive interventions are impossible. As a paradigmatic type of pathological propagation, we considered seizure spread. Our in-silico approach allowed an exhaustive search, which would have been impossible in vivo. The model predicts the activity propagation of a seizure originating in a certain epileptogenic zone and to identify the most unstable pathways that support and allow the propagation. In other words, activity propagation cannot be disjointed by the network topology outside the epileptogenic zone and by the topological features linked to the propagation zone that both contribute to the propagation. We have demonstrated that the energy dissipation of seizure propagation may be controlled by performing a Linear Stability Analysis of the system, thus allowing identification of the properties and the response of the system undergoing a perturbation (such as a seizure emission). In this framework, the most unstable directions that carry on the spreading may be identified and, by studying the effect of lesions on the dynamics of the simulated brain processes, it is possible to visualize when the seizure stops propagating, and the recruitment of the PZ does not occur anymore. A similar stability analysis has been performed in [39] on a functional connectivity matrix reconstructed from ECoG recordings, where finite perturbations applied to single nodes render the stable network unstable causing a seizure. The identification of the most fragile nodes, able to desynchronize the network, turns out to be fundamental to construct a predicted EZ set. This complementary study reinforce the hypothesis that dynamics matter and the nature of seizure onset and spreading cannot be captured by the structural connectivity alone as we have shown here controlling the propagation via the Linear Stability Analysis: the Jacobian matrix of the system takes into account not only the underlying structural network but also the dynamical variables of the system.

Other groups have already studied the effects of removing brain regions and connections on the dynamics of simulated network processes [40], but in all these works the interest is mainly focused on understanding the effect of damages on the brain activity, instead of using selected lesions as the least invasive treatment as possible in an eventual surgical operation. Moreover, our approach is entirely based on structural data, which allows the creation of a brain model based on purely non-invasive data prior to any surgery or exploration via intracranial measurements, whereas other techniques rely on invasive functional data focused on the estimation of the epileptogenic zone [41] and resective surgery [42, 43]. Disconnection approaches thus are significantly less invasive and bear the potential for general network control and applications beyond seizure suppression.

Our approach generalizes to oscillatory activity propagation so far, that it does not necessitate the presence of the hallmark of epilepsy, that is an onset and offset of high-frequency oscillatory activity. It thus can be applied in the future to understand the directionality of state changes and manipulate the activity in the network. On the other hand, concerning the applicability to the seizure propagation problem, it may be determinant to design a new surgical approach, providing support and improving the resection surgical procedure. A less invasive and more effective surgical approach would make the difference in short and longterm outcomes and in improving the quality of life of the patients. In particular reducing the involvement of propagation networks is a major factor to reduce the impact of seizures, particularly the loss of consciousness [44], which is one of the major seizure signs and it is clearly linked to the synchronization in propagation network, specially fronto-parietal networks in temporal lobe epilepsy. Therefore by blocking the seizure propagation we can improve the quality of life of epileptic patients, although we recognize that seizure freedom with a classification IA according to Engel is the real goal of epilepsy surgery. Seizure free patients classified IB according to the Engel may include patients with residual subjective symptoms (aura) but without objective signs (automatism, loss of consciousness, etc.). Moreover, an eventual persistence of seizures after surgery would not inhibit a second clinical trial for patients, since the procedure does not consist in removing consistent parts of the brain, that on the contrary, would impair the physical abilities of the patient.

## V. METHODS

### A. Patient selection and data acquisition

We selected 15 drug-resistant patients (9 females, 6 males, mean age 33.4, range 22-56) with different types of partial epilepsy accounting for different EZ localizations. All patients underwent a presurgical evaluation (see Supplementary Table 1). For each patient a first not invasive evaluation procedure is foreseen: this evaluation comprises of the patient clinical record, neurological examinations, positron emission tomography (PET), and electroencephalography (EEG) along with video monitoring. T1 weighted anatomical images (MPRAGE sequence, TR=1900 ms, TE=2.19 ms, 1.0 x 1.0 x 1.0 mm, 208 slices) and diffusion MRI images (DTI-MR sequence, angular gradient set of 64 directions, TR=10.7 s, TE=95 ms, 2.0 x 2.0 x2.0 mm, 70 slices, b weighting of 1000 *s/mm^2^*) were acquired on a Siemens Magnetom Verio 3T MR-scanner. From the gathered data clinicians conclude potential epileptogenic zones (EZ). Further elaboration on the EZ are done in the second phase, which is invasive and consists of placement of stereotactic EEG (SEEG) electrodes in or close to the suspected regions. These electrodes are equipped with 10 to 15 contacts that are 1.5 mm apart. Each contact is 2 mm of length and 0.8 mm in diameter. The SEEG was recorded by a 128 channel Deltamed™ system using a 256 Hz sampling rate. The SEEG recordings were band-pass filtered between 0.16 and 97 Hz by a hardware filter. All the chosen patients showed seizures in the SEEG starting in one or several localized areas (EZ), before recruiting distant regions (PZ). The position of the electrodes was pinned down by performing a computerized tomography (CT) scan or a MRI after implanting the electrodes.

### B. Data processing

The data processing to import structural and diffusion MRI data in The Virtual Brain has been done using SCRIPTS. This processing pipeline makes use of various tools such as FreeSurfer [45], FSL [46], MRtrix3 [47] and Remesher [48], to reconstruct the individual cortical surface and large-scale connectivity. The surface was reconstructed using 20,000 vertices. Cortical and volumetric parcellations were performed using the Desikan-Killiany atlas with 70 cortical regions and 17 subcortical regions [49] (one more empty region is added in the construction of the structural connectivity for symmetry). Two additional parcellations were obtained by subdividing each cortical region of the Desikan-Killiany atlas in two and four: it has been possible to obtain parcellations with 157 and 297 regions, respectively [50]. Numerical simulations performed with different parcellations show that the large-scale network dynamics does not change significantly, while the local dynamics becomes richer with increasing parcellation size. We here use a parcellation size that has been judged reasonable by Proix et al. [51] to address network effects.

The diffusion data was corrected for eddy-currents and head motions using eddy-correct FSL functions. Fiber orientation estimation was performed with Constrained Spherical Deconvolution [52], and improved with Anatomically Constrained Tractography [53]. Tractography was performed using 2.5 10^6^ fibers and it was corrected using Spherical-Deconvolution Informed Filtering of Tractograms [54]. The connectivity matrix was obtained by summing track counts over each region of the parcellation; when performing the numerical analysis the matrix had been normalized so that the maximum value of the connectivity matrix was one.

The CT or MRI scan performed after electrode placements were aligned with the structural MRI recorded before the surgery using the FLIRT function of FSL, with 6 degrees of freedom and a Mutual Information cost function. Each contact surface was reconstructed and assigned to the region of the corresponding parcellations containing most of the contact volume.

### C. Clinical definition of the Propagation Zone

The propagation was defined by two different methods. The first method is the subjective evaluation of clinicians based on the different measurement modalities (EEG and SEEG) gathered throughout the two-step procedure (non-invasive and invasive). The second method is objective in the sense that it is solely based on the SEEG measurements. For each patient, all seizures were isolated in the SEEG time series. The bipolar SEEG was considered (between pairs of electrode contacts) and filtered between 1-50 Hz using a Butterworth bandpass filter. A contact was considered to be in the PZ if its signal energy was responsible for at least 30% of the maximum signal energy over the contacts, and was not in the EZ. The corresponding region was then assigned to the PZ.

### D. The Model

Large scale brain network activity and recruitment effects shown previously have been obtained by simulating the dynamical evolution of a phenomenological dynamic model called Epileptor. In particular the dynamics of the single node coupled in a network of N elements read

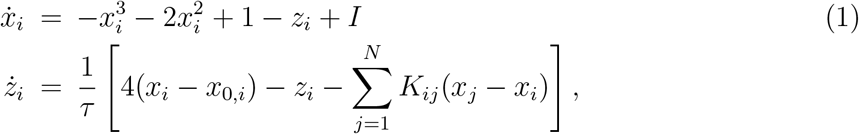

with *τ* = 2857 and *I* = 3.1 passive current setting the operating point of the Epileptor. Eqs. 1 are derived from a more complex 5-dim model which takes into account the timescale separation present in the seizure evolution (the derivation of eqs. 1 from the complete 5-dim. model is reported in the SI, see Text S1). Here the timescale difference is guaranteed by *τ* ≫ 1. On the fastest timescale the state variable *x_i_* exhibits bistable dynamics between oscillatory activity, thus modeling ictal activity, and a stable node representing the healthy condition. On the slowest timescale, the evolution of the permittivity variable *z_i_* guides the neural population through the seizures, capturing the details of the autonomous slow evolution of interictal and ictal phases. Moreover the coupling function 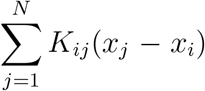 expressed as a linear difference coupling term, models distant discharges at location *j* as first order deviations from the homeostatic equilibrium of the slow permittivity variable *z_i_*. *K* is the structural connectivity matrix whose entries *K_i,j_*· are real numbers rescaled with the maximal entry value; they represent the connections between the brain populations *i* and *j*. In particular the structural connectivity matrix of each single patient is described in terms of 88 populations given the parcellation of the brain into 88 main regions. Finally *x_0,i_* represents the degree of epileptogenicity and its value is chosen according to the relationship Δ*x*_0,*i*_ = *x*_0,*i*_ – *x_c_*, with *x_c_* = –2.1 being the critical value between the stable (epileptogenic) and unstable (healthy) regime. If Δ*x*_0,*i*_ > 0, a brain region is epileptogenic and seizures are triggered autonomously. Otherwise, if Δ*x*_0,*i*_ < 0 regions are in an equilibrium state. In our simulations only the epileptogenic zone, selected according to medical predictions, has a degree of epileptogenicity such that Δ*x*_0,*i*_ > 0; for all the other regions Δ*x_0,i_* < 0.

### E. Linear stability analysis: eigenvalues and eigenvectors

The stability of eqs. 1 can be analyzed by following the evolution of infinitesimal perturbations in the tangent space, whose dynamics is ruled by the linearization of eqs. 1 as follows

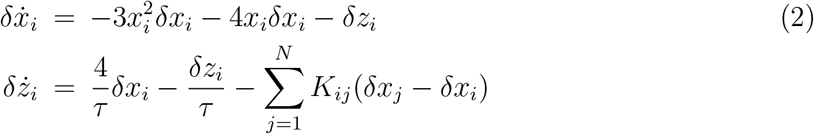

In particular, if we denote with 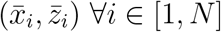 the steady state solution of eqs. 1, the stability of this state is determined by the eigenvalues of the Jacobian matrix J calculated in 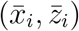, where *J* is given by

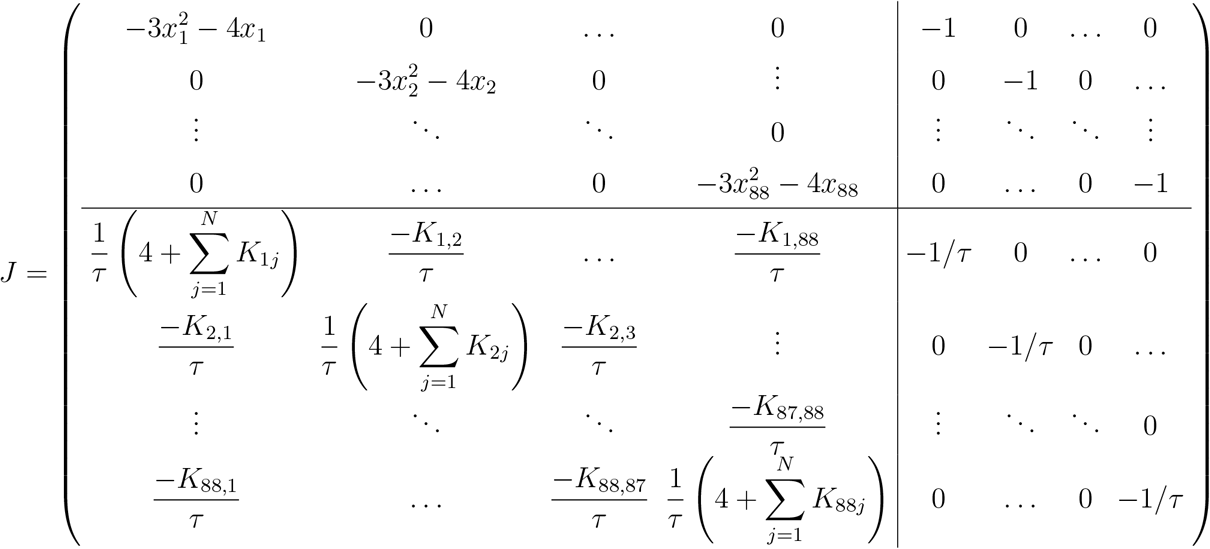

If the eigenvalues of the Jacobian matrix all have real parts less than zero, then the steady state is stable. In terms of seizure propagation, this means that the seizure should not be able to propagate. On the other hand, if at least one of the eigenvalues of the Jacobian matrix has real part greater than zero, then the steady state is unstable. In this second case we expect a seizure to propagate along the network.

Moreover the eigenvectors of the Jacobian matrix give the axes along which the behaviors indicated by the eigenvalues are centered. This means that, in case of negative eigenvalue, the associated eigenvector points in the direction along which the fixed point attracts. In case of positive eigenvalue, the eigenvector points the direction along which the fixed point repels. Therefore the computation of the eigenvectors allow us to identify the pathways along which the seizure propagates. In particular if we identify the largest components of the maximal eigenvector, i.e. the largest elements of the eigenvector related to the maximal (positive) eigenvalue, we can straightforwardly suggest the most unstable directions able to convey the propagation along the entire network. In other words the elements corresponding to the largest components of the maximal eigenvector will be the most easily recruited during the propagation. On this basis we estimated the Propagation Zone (PZ) by identifying the elements with the biggest absolute value of the maximal eigenvector: these elements represent the dominating sub-networks involved in the transition towards the seizure state. These results can be compared with the PZ which is empirically obtained by neurosurgeons or during the data analysis of the time series of the cortical and implanted EEG electrodes [55], resulting in a good agreement.

Finally it is worth noticing that in the Jacobian matrix J, appear not only the structural connectivity matrix *K_ij_* but also the dynamical variables of the system: the recruitment process is a dynamical one and cannot be predicted by the connectivity alone. Therefore a combination of structural and dynamical information as it is used in the Linear Stability Analysis, is fundamental to predict the PZ and to suggest minimally invasive disconnections.

### F. Prospects for personalized spatial propagation zones

The hypothesis of EZ, estimated by medical doctors, is implemented in the model eqs. 1 and in the analysis, by two different levels of excitability *x*_0_: *x*_0_ < *x_c_* for the epileptic zone nodes, *x*_0_ > *x_c_* for the healthy nodes. There is no need to define a third level of excitability for the nodes in the propagation zone to describe the recruitment mechanism, since PZ emerges self-consistently from the dynamics. Furthermore, the Jacobian matrix is an explicit function of the fixed points of the model, meaning that the eigenvectors and eigenvalues obtained from the Linear Stability Analysis will also be influenced.

The SEEG samples neuroelectric activity from various sources, which are more than the number of contacts on the electrods. This undersampling would pose an enormous problem if we were inverting the SEEG signals to source signals, which is an ill-posed problem. Instead we compute the physical forward solution and then perform validation in signal space (rather than source space). Therefore, even though contacts are limited in number and in placement positions, our capacity of model identification is not affected by undersampling.

### G. Network measures

Topological properties of a network can be examined by using different graph measures that are provided by the general framework of the graph theory. These graph metrics can be classified in measures that are covering three main aspects of the topology: segregation, integration and centrality. The segregation accounts for the specialized processes that occur inside a restricted group of brain regions, usually densely connected. Among the possible measures of segregation, we have considered the *clustering coefficient,* which gives the fraction of triangles around a node and it is equivalent to the fraction of node’s neighbors that are neighbors of each other as well. This is an important measure of segregation, since the presence of a dense neighborhood around a node is fundamental to the generation of clusters and cliques that are likely to share specialized information.

The measures of integration refer to the capacity of the network to rapidly combine specialized information from not nearby, distributed regions. Integration measures are based on the concept of communication paths and path lengths, which estimate the unique sequence of nodes and links that are able to carry the transmission flow of information between pairs of brain regions. In particular the path length measures the sum of the edge weights in a weighted graph (as the one that we have considered) and the shortest path length suggest a stronger potential for integration between different brain regions. At the global level it is moreover possible to define a “characteristic path length” of the network, which is calculated as the average inverse shortest path length between the all pairs of nodes in the network. Among the possible measures of integration, as the most meaningful for our structural connectivity matrices we have chosen the *efficiency.* It is defined as the the average inverse shortest path length in the network, and is inversely related to the characteristic path length.

Centrality refers to the importance of network nodes and edges for the network functioning. The most intuitive index of centrality is the *node degree,* which gives the number of links connected to the node; for this measure connection weights are ignored in calculations. The *node strength* is the sum of weights of links connected to the node, while node *betweenness centrality* is the fraction of all shortest paths in the network that contain a given node. Nodes with high values of betweenness centrality participate in a large number of shortest paths. Finally, *closeness centrality* is the inverse of the average shortest path length from one node to all other nodes in the network.

## Acknowledgments

S.0. acknowledge Benjamin Lindner, Dionysios Perdikis, Timothée Proix, Marmaduke Woodman for useful discussions. This work was partially supported by The Short Term Mobility Programm founded by the Italian National Research Council (protocol number 0057676, date 31/08/2015) and by the Project Epinext – Federation Hospitalo-Universitaire (FHU) “Epilepsy and Disorders of Neuronal Excitability”.

## Author Contributions

Conceived and designed the experiments: SO VJ. Performed the experiments and analyzed the data: SO SP. Provided the data: MG FB. Wrote the paper: SO SP MG FB VJ

## Additional Information

Competing financial interests: The authors declare no competing financial interests. Non-financial competing interests: The authors declare no competing non-financial interests.

## Supporting Information

**Text S1: Model – The model**

**Text S2: Data – Topological properties**

**Text S3: Numerical simulations – Other patients**

**Figure S1:**

**Model – Comparison between the maximum Lyapunov vector and the maximum eigenvector.** Panel a: Localization of the maximum Lyapunov vector, correspondent to the maximum Lyapunov exponent. Panel b: Maximum eigenvector as a function of the node index. The dashed lines indicate the node that are first recruited in the propagation zone and whose links with the epileptogenic zone represent the directions along which the seizure spreads. For both panels the structural connectivity matrix of patient CJ has been used.

**Figure S2:**

**Data – Topological properties of patient’s connectome**. Left panel: Efficiency value of each population. Right panel: Strength value of each population. Both properties are calculated on the structural connectivity matrix of the patient CJ. The clinical estimated EZ is highlighted in yellow, while the estimated PZ (predicted via Linear Stability Analysis) is highlighted in magenta.

**Figure S3:**

**Data – Topological properties of patient’s connectome**. Left panel: Clustering value of each population. Right panel: Degree value of each population. Both properties are calculated on the structural connectivity matrix of the patient CJ. The clinical estimated EZ is highlighted in yellow, while the estimated PZ (predicted via Linear Stability Analysis) is highlighted in magenta.

**Figure S4:**

**Data – Topological properties of patient’s connectome**. Left panel: Betweeness value of each population. Right panel: Centrality value of each population. Both properties are calculated on the structural connectivity matrix of the patient CJ. The clinical estimated EZ is highlighted in yellow, while the estimated PZ (predicted via Linear Stability Analysis) is highlighted in magenta.

**Figure S5:**

**Numerical simulations – Analysis of patient FB.** Left panel: Seizure events as a function of time. EZ (red dots): rh-Precentral Gyrus that corresponds to node 77. Disconnections PrG-PoG, PrG-CMFG, PrG-PoP, PrG-SFG, PrG-Th, PrG-Pu, PrG-PaC, PrG-SMG must be performed in order to stop the seizure. All the disconnections are performed among populations belonging to the right hemisphere. Dashed blue line corresponds to the time at which lesions occur. Right panel: Number of lesions necessary to stop seizure propagation as a function of the epileptogenic region.

**Figure S6:**

**Numerical simulations – Analysis of patient ET.** Left panel: Seizure events as a function of time. EZ (red dots): lh-Posterior Cingulate Gyrus, lh-Precuneus Cortex corre sponding respectively to nodes 33 and 35. Partial targeted disconnection procedure to be performed in order to stop seizure propagation: PCunC-SPC, PCunC-PCG, PCunC-ICC, PCunC-Cun, PCunC-rPCunC, PCunC-Pac, and the links between PCG and nodes SPC, ICC, Cun, rPCunC, Pac, SFG, CACC. The targeted disconnection is performed among populations in the left hemisphere. Dashed blue line corresponds to the time at which lesions occur. Right panel: Number of lesions necessary to stop seizure propagation as a function of the epileptogenic region.

**Figure S7:**

**Numerical simulations – Analysis of patient AC.** Left panel: Seizure events as a function of time. EZ (red dots): rh-Lateral Orbito Frontal Cortex, rh-Temporal pole corresponding respectively to nodes 65 and 86. Partial targeted disconnection procedure to be performed in order to stop seizure propagation: LOFC-RMFG. The targeted disconnection is performed among populations in the right hemisphere. Dashed blue line corresponds to the time at which lesions occur. Right panel: Number of lesions necessary to stop seizure propagation as a function of the epileptogenic region.

**Figure S8:**

**Numerical simulations – Analysis of patient GC.** Left panel: Seizure events as a function of time. EZ (red dots): rh-Hippocampus, rh-Amygdala corresponding respectively to nodes 51 and 52. Partial targeted disconnection procedure to be performed in order to stop seizure propagation: Hi-EntC, Hi-PHiG, Hi-Pal, Hi-Th, Hi-FuG, Hi-Amg, Amg-TmP. The targeted disconnection is performed among populations in the right hemisphere. Dashed blue line corresponds to the time at which lesions occur. Right panel: Number of lesions necessary to stop seizure propagation as a function of the epileptogenic region.

**Figure S9:**

**Numerical simulations – Analysis of patient CV.** Left panel: Seizure events as a function of time. EZ regions (red dots): Posterior Cingulate Gyrus (node 33), Caudal

Middle Frontal Gyrus (node 14), Superior Frontal Gyrus (node 38). All the areas belonging to the EZ are located in the left hemisphere. Links between regions SFG-RMFG, SFG-(rh-SFG) and PCG-CMFG are cut in order to stop the seizure. Dashed blue line corresponds to the time at which lesions occur. Right panel: Number of lesions necessary to stop seizure propagation as a function of the epileptogenic region.

**Figure S10:**

**Numerical simulations – Analysis of patient RB.** Left panel: Seizure events as a function of time. EZ regions (red dots): Amygdala (node 9), Hippocampus (node 8), Entorhinal cortex (node 16), Fusiform gyrus (node 17), Temporal Pole (node 43), rh-Entorhinal cortex (node 59). All the areas belonging to the EZ are located in the left hemisphere. Links between all the regions in the EZ and the regions in the PZ must be cut in order to stop the seizure. Dashed blue line corresponds to the time at which lesions occur. Right panel: Number of lesions necessary to stop seizure propagation as a function of the epileptogenic region.

**Table S1:**

**Data – Clinical characteristics of each patient.** Partial seizures were recorded with stereotactic EEG (SEEG) electrodes in 15 drug-resistant epileptic patients undergoing presurgical evaluation. The clinical characteristics of each patient are given in Table ??: N, normal; L, left; R, right; Th, thermocoagulation; Gk, Gamma knife; Sr, surgical resection; NO, not operated; PVH, periventricular nodular heterotopia; FCD, focal cortical dysplasia; SPC, superior parietal cortex; F, Frontal; NA, not available.

