## Supplementary Information for "Controlling seizure propagation in large-scale brain networks"

### Text S1: The Model

#### A. The 5-dim Epileptor model

In order to take into account the timescale separation present in the seizure evolution, Jirsa et al. [1] used the theory of fast-slow systems in nonlinear dynamics [2] to develop a taxonomy of seizures. In addition, from experimental seizure data of various species, they identified a predominant class of bifurcation pairs, which resulted to be integrated into a phenomenological dynamic model called Epileptor. The Epileptor is a five-dimensional model and comprises three different timescales accounting for various electrographic patterns: on the fastest timescale, two state variables ($x_1$ and $y_1$) exhibit bistable dynamics between oscillatory activity, thus modeling fast discharges, and a stable node representing interictal activity. On the intermediate timescale two state variables ($x_2$ and $y_2$) model the spike and wave events. Finally, on the slowest timescale, the evolution of a very slow permittivity variable ($z$) guides the neural population through the seizures, including seizure onset and offset. In particular the permittivity variable captures the details of the autonomous slow evolution of interictal and ictal phases, as well as various details of seizure evolution during each phase. The activity in this model is autonomously switching between interictal and ictal states because of the slow permittivity variable accounting for the extracellular effects related to energy consumption and tissue oxygenation. In particular the activity is described by the following equations

$$\begin{aligned}
&\dot{x}_1 = y_1 - f_1(x_1, x_2) - z + I_1, \qquad \dot{y}_1 = 1 - 5x_1^2 - y_1 \\
&\dot{z} = \frac{1}{\tau_0}\left[4(x_1 - x_0) - z\right] \\
&\dot{x}_2 = -y_2 + x_2 - x_2^3 + I_2 + 0.002g(x_1) - 0.3(z - 3.5), \qquad \dot{y}_2 = \frac{1}{\tau_2}(-y_2 + f_2(x_1, x_2)),
\end{aligned} \tag{1}$$

where

$$f_1(x_1, x_2) = \begin{cases} x_1^3 - 3x_1^2 & \text{if } x_1 < 0; \\ (x_2 - 0.6(z-4)^2)x_1 & \text{if } x_1 \geq 0. \end{cases}$$

$$f_2(x_1, x_2) = \begin{cases} 0 & \text{if } x_2 < -0.25; \\ 6(x_2 + 0.25)x_1 & \text{if } x_1 \geq -0.25. \end{cases}$$

$$g(x_1) = \int_{t_0}^{t} e^{-\gamma(t-\tau)} x_1(\tau) d\tau.$$

and the degree of epileptogenicity is represented by the value $x_0$. If we identify with $x_c$ the critical value between the stable and unstable regime, for $x_0 < x_c$, the Epileptor autonomously shows seizure activity and is said to be epileptogenic while, for values of $x_0 > x_c$, the Epileptor is in its (healthy) equilibrium state. It is worth noticing that the distinction between excitation and inhibition in terms of explicit mechanistic realizations (for instance, GABA, Glutamate, etc.) is impossible in the model. These mechanisms are absorbed in the generic dynamics of the Epileptor model. The distinction of excitation and inhibition is, however, still preserved functionally, that is in terms of their role in the model linked to increase or decrease of state variables values. The functional distinction (as opposed to mechanistic) of excitation/inhibition persists through the mathematical expression of bifurcations in terms of the capability of generating oscillatory behavior (corresponding to the epileptogenic state) or not. A detailed mechanistic realization of the Epileptor model has been given in [3] where a correspondence between the emergent dynamic behaviors of this model and a network model of spiking neurons is investigated. In particular Naze *et al.* consider two neuronal populations, characterized by fast excitatory bursting neurons and regular spiking inhibitory neurons, embedded in a common extracellular environment represented by a slow variable: in this setup spike waves, including interictal spikes, are generated primarily by inhibitory neurons, whereas fast discharges during the wave part are due to excitatory neurons. The system is pushed into paroxysmal regimes by slow variations of global excitability, due to exogenous fluctuations from extracellular environment, and by gap junction communication. However the mechanistic realization shown in [3] is not unique and depends on the chosen spiking network model. It can be, for instance, easily demonstrated analytically that the system's oscillation amplitude depends only on the E/I balance and cannot isolate the individual contributions. There is an extensive general literature on the identifiability of model mechanisms, whose practical use is unfortunately limited to a small number of degrees of freedom [4]. Equations 1 describe the dynamics of a single, uncoupled phenomenological model of epileptiform neural activity. Using again time-scale separation as well as evidences from SEEG recordings, Proix *et al.* [5] introduced a nonlocal interareal permittive coupling between Epileptors and they suggested to consider seizure recruitment among brain regions on a slow time scale. Such a coupling function can be expressed as a linear difference coupling term, assuming first order deviations from the homeostatic equilibrium of the slow permittivity variable. In particular it is possible to approximate the effect of fast neuronal discharges as a homeostatic perturbation of the slow permittive variable $z$ of the Epileptor in the target region. Therefore the distant discharges at remote location $j$ are transmitted via axons to target region $i$ and can be seen as a perturbation of the homeostatic equilibrium of the region $i$. This disturbance of ion homeostasis will depend on both local and distant neuronal discharges and can be schematized as a coupling term $C_{i,j}(x_{1,i}, x_{1,j}) = \frac{1}{\tau_0}\sum_{j=1}^{N} k_{ij}(x_{1,j} - x_{1,i})$ that influences the rate of change of the permittivity variable $z_1$ of the region $i$, for $i, j = 1, ..., N$, where $N$ is the number of regions of interest. Under these considerations, the final model equations for N-coupled Epileptors with permittive coupling read as follows:

$$
\begin{aligned}
&\dot{x}_{1,i} = y_{1,i} - f_1(x_{1,i}, x_{2,i}) - z_i + I_1, \qquad \dot{y}_{1,i} = 1 - 5x_{1,i}^2 - y_{1,i} \\
&\dot{z}_i = \frac{1}{\tau_0}\left[4(x_{1,i} - x_{0,i}) - z_i\right] - \frac{1}{\tau_0}\sum_{j=1}^{N} k_{ij}(x_{1,j} - x_{1,i}) \\
&\dot{x}_{2,i} = -y_{2,i} + x_{2,i} - x_{2,i}^3 + I_2 + 0.002g(x_{1,i}) - 0.3(z_i - 3.5), \quad \dot{y}_{2,i} = \frac{1}{\tau_2}(-y_{2,i} + f_2(x_{1,i}, x_{2,i})),
\end{aligned}
\tag{2}
$$

where $\tau_0 = 2857$, $\tau_2 = 10$, $I_1 = 3.1$ and $I_2 = 0.45$; $f_1$, $g$ and $f_2$ are defined as above.

#### B. Two dimensional reduction of the Epileptor

To study the recruitment effects on the slowest timescale of the permittivity variable, and in particular to investigate large scale brain networks, it is possible to apply averaging methods to the coupled Epileptor equations, in order to reduce the dimension of the original model. Numerical analyses of the full system demonstrate that the effect of the second neuronal ensemble ($x_{2,i}$ and $y_{2,i}$) is negligible under this approximation, hence we can neglect it. Then the Epileptor equations (2) become

$$\dot{x}_{1,i} = -x_{1,i}^3 - 2x_{1,i}^2 + 1 - z_i + I_i \tag{3}$$
$$\dot{z}_i = \frac{1}{\tau_0}\left[4(x_{1,i} - x_{0,i}) - z_i - \sum_{j=1}^{N} K_{i,j}(x_{1,j} - x_{1,i})\right].$$

For the sake of simplicity of our analysis, we also drop the subscripts $_1$ in the previous equations. Finally, the model used to mimic the dynamics of the single node coupled in a network reads

$$\dot{x}_i = -x_i^3 - 2x_i^2 + 1 - z_i + I \tag{4}$$
$$\dot{z}_i = \frac{1}{\tau}\left[4(x_i - x_{0,i}) - z_i - \sum_{j=1}^{N} K_{ij}(x_j - x_i)\right],$$

with $\tau = 2857$, $I = 3.1$. The timescale difference is guaranteed by $\tau \gg 1$. Even in the reduced model $x_{0,i}$ represents the degree of epileptogenicity and its value is chosen according to the relationship $\Delta x_{0,i} = x_{0,i} - x_c$, with $x_c = -2.1$ being the critical value between a stable and an epileptogenic Epileptor. If $\Delta x_{0,i} > 0$, a brain region is epileptogenic and seizures are triggered autonomously. Otherwise, $\Delta x_{0,i} < 0$ and regions are in an equilibrium state. $K_{i,j}$ represents the coupling matrix between the brain populations; the structural connectivity matrix of single patients is used with 88 populations describing the main regions in the brain. The entries in the matrix are rescaled with the value of the maximal entry.

#### C. Linear stability analysis: eigenvectors vs Lyapunov vector

From a numerical point of view the fixed point solution $(\bar{x}_i, \bar{z}_i)$ $\forall i \in [1, N]$ of the system can be found by setting $\dot{x}_i = 0, \dot{z}_i = 0$ and by implementing the Newton's function, a one-dimensional root-finding routine which is also called the Newton-Raphson method. While all the healthy nodes are subject to a constant motion in $x_i$, the epileptogenic node shows a periodic motion and, a priori, its stable state cannot be restricted to a simple fixed point solution in order to calculate the Linear Stability Analysis around it. However this operation is justified by the time scale separation. In particular it is always possible to compute the stability around the stable manifold: this is called dissection analysis and holds if two separate time scales are present [6]. Once the fixed point is found, its stability is determined by calculating both the eigenvalues and eigenvectors of the Jacobian matrix. The Jacobian matrix J is calculated according to the following system:

$$\begin{aligned} \delta\dot{x}_i &= -3x_i^2\delta x_i - 4x_i\delta x_i - \delta z_i \\ \delta\dot{z}_i &= \frac{4}{\tau}\delta x_i - \frac{\delta z_i}{\tau} - \sum_{j=1}^{N} K_{ij}(\delta x_j - \delta x_i) \end{aligned} \tag{5}$$

and it is given by

$$J = \left(\begin{array}{cccc|cccc} -3x_1^2 - 4x_1 & 0 & \dots & 0 & -1 & 0 & \dots & 0 \\ 0 & -3x_2^2 - 4x_2 & 0 & \vdots & 0 & -1 & 0 & \dots \\ \vdots & \ddots & \ddots & 0 & \vdots & \ddots & \ddots & \vdots \\ 0 & \dots & 0 & -3x_{88}^2 - 4x_{88} & 0 & \dots & 0 & -1 \\ \hline \frac{1}{\tau}\left(4 + \sum_{j=1}^{N} K_{1j}\right) & \frac{-K_{1,2}}{\tau} & \dots & \frac{-K_{1,88}}{\tau} & -1/\tau & 0 & \dots & 0 \\ \frac{-K_{2,1}}{\tau} & \frac{1}{\tau}\left(4 + \sum_{j=1}^{N} K_{2j}\right) & \frac{-K_{2,3}}{\tau} & \vdots & 0 & -1/\tau & 0 & \dots \\ \vdots & \ddots & \ddots & \frac{-K_{87,88}}{\tau} & \vdots & \ddots & \ddots & 0 \\ \frac{-K_{88,1}}{\tau} & \dots & \frac{-K_{88,87}}{\tau} & \frac{1}{\tau}\left(4 + \sum_{j=1}^{N} K_{88j}\right) & 0 & \dots & 0 & -1/\tau \end{array}\right)$$

The eigenvalues indicate the stability of the system (4): positive eigenvalues correspond to an unstable system and in this case we expect a seizure to propagate along the network. On the other hand if the eigenvalues are non-positive the system is stable and the seizure should not be able to propagate.

Moreover the computation of the eigenvectors allow us to identify the pathways along which the seizure propagates. In particular it is possible to identify the biggest components of the maximal eigenvector, i.e. the eigenvector related to the maximal eigenvalue, as the most unstable directions able to convey the propagation in the entire network. In other words the elements with the biggest components of the maximal eigenvector will be the most probable to be recruited during the propagation. On this basis we estimated the Propagation Zone (PZ) by identifying the elements with the biggest amplitude: these elements represent the dominating sub-networks involved in the transition towards the seizure state. These results can be compared with the PZ which is empirically obtained by neurosurgeons or during the data analysis of the time series of the cortical and implanted EEG electrodes [7].

In order to support our analysis of the most unstable directions identified via the Linear Stability Analysis, we have compared the prediction obtained via the maximum eigenvector with the localization of the maximum Lyapunov vector. Lyapunov vectors describe the characteristic expanding and contracting directions of a dynamical system. These vectors are defined along the trajectories of a dynamical system: in particular, if the system can be described in terms of a $n$-dimensional state vector, $\mathbf{x} \in R^n$, the Lyapunov vectors $\mathbf{v}^k(x)$, $(k = 1 \ldots n)$ point in the directions in which an infinitesimal perturbation will grow asymptotically, exponentially at an average rate given by the Lyapunov exponents. When expanded in terms of Lyapunov vectors, a perturbation asymptotically aligns with $\mathbf{v}^1$, i. e. the Lyapunov vector in that expansion corresponding to the largest Lyapunov exponent, as this direction outgrows all others. Therefore almost all perturbations align asymptotically with the Lyapunov vector corresponding to the largest Lyapunov exponent in the system. In order to numerically calculate $\mathbf{v}^1(x(t))$, we computed the exponential growth rate of the maximum Lyapunov exponents along the 2N-1 independent directions in tangent space. The Lyapunov exponents has been numerically estimated by implementing the standard algorithm [8] on the time-dependent set of eqs. (5). Finally the localization $\mathcal{L}$ of the maximal Lyapunov vector is obtained averaging in time the square modulus of the vector itself: $\mathcal{L} =< \sum_{i=1}^{n} v_i^1(t) * v_i^1(t) >_t$.

In this system the Lyapunov spectrum does not show any level of chaoticity (the exponents are usually non-positive) and the localization of the maximal Lyapunov vector allows us to understand the origin of the instability responsible for the seizure spreading, thus making possible to detect the direction of maximum instability of the system. The directions of maximum instability coming out from the localization analysis are the same as the ones previously calculated via the Linear Stability Analysis and reported in Fig. 5(b). The comparison between the two analyses is shown in Fig. S1, where both the localization of $\mathbf{v}^1$ and the maximum eigenvector are reported; in particular the dashed lines highlight the common maximum elements, that represent the directions along which the seizure spreads.

A priori this is not obvious since Lyapunov vectors are not usually identical with the local principal expanding and contracting directions, i.e. the eigenvectors of the Jacobian. In fact, while the latter require only local knowledge of the system, the Lyapunov vectors are influenced by all Jacobians along a trajectory, thus representing a measure of the instability of the dynamics of the system, not related to any particular fixed point or initial stable solution.

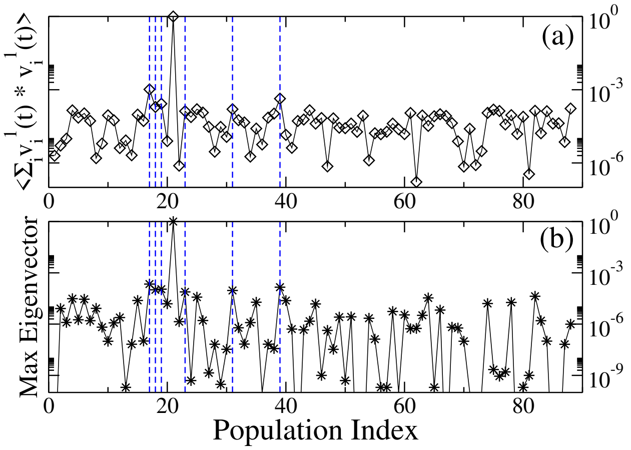


Figure 1: Panel a: Localization of the maximum Lyapunov vector, correspondent to the maximum Lyapunov exponent. Panel b: Maximum eigenvector as a function of the node index. The dashed lines indicate the node that are first recruited in the propagation zone and whose links with the epileptogenic zone represent the directions along which the seizure spreads. For both panels the structural connectivity matrix of patient CJ has been used.

#### D. Prospects for personalized spatial propagation zones

The knowledge of PZ and EZ, estimated by medical doctors, can be easily implemented in the model eqs. (1, 2) and in the analysis, by simply defining three different levels of excitability $x_0$: $x_0 < x_c$ for the epileptic zone nodes, $x_0 \gg x_c$ for the healthy nodes, and $x_0 > x_c$, where $x_0 - x_c \ll 0$ for the healthy nodes in the propagation zone where the epileptic activity can spread. On the other hand, only two different levels of the degree of epileptogenicity $x_0$ are necessary to enter in eq. (5), thus directly affecting its solutions, i. e. fixed points $(\bar{x}_i, \bar{z}_i)$ $\forall i \in [1, N]$. In particular for the reduced system it is not necessary to take a different level of epileptogenicity for the nodes in the propagation zone to describe the recruitment mechanism. Furthermore, the Jacobian matrix is an explicit function of the fixed points of the reduced epileptor, meaning that the eigenvector and eigenvalues obtained from the Linear Stability Analysis will also be influenced. Henceforth, a similar comparison like above can be performed for the more realistic scenarios that involve the PZ, and the outcomes of different lesioning strategies can be also computed and compared.

### I. TOPOLOGICAL PROPERTIES

Topological properties of the structural connectivity matrix of patient **CJ**, examined by means of 6 different graph metrics provided by the general framework of graph theory. These metrics are Efficiency (Fig. **??**), Strength (Fig. **??**), Clustering (Fig. **??**), Degree (Fig. **??**), Betweeness (Fig. **??**), Centrality (Fig. **??**). Neither the epileptogenic zone (highlighted in each panel with a yellow stripe) nor the propagation zone (highlighted in magenta) are characterized by peculiar values of these metrics, thus suggesting that network measures cannot be used to identify biomarkers of these specific brain functions.

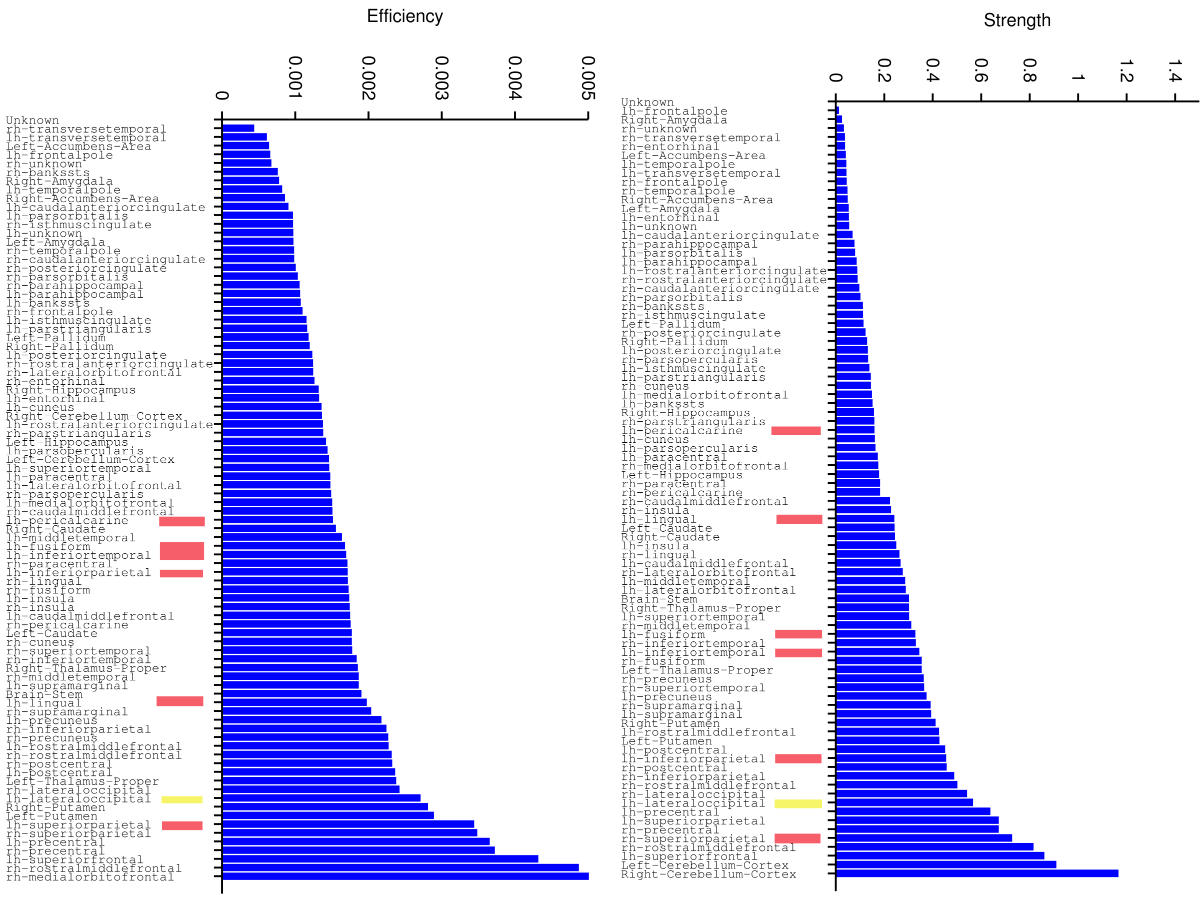


Figure 2: Left panel: Efficiency value of each population. Right panel: Strength value of each population. Both properties are calculated on the structural connectivity matrix of the patient **CJ**. The clinical estimated EZ is highlighted in yellow, while the estimated PZ (predicted via Linear Stability Analysis) is highlighted in magenta.

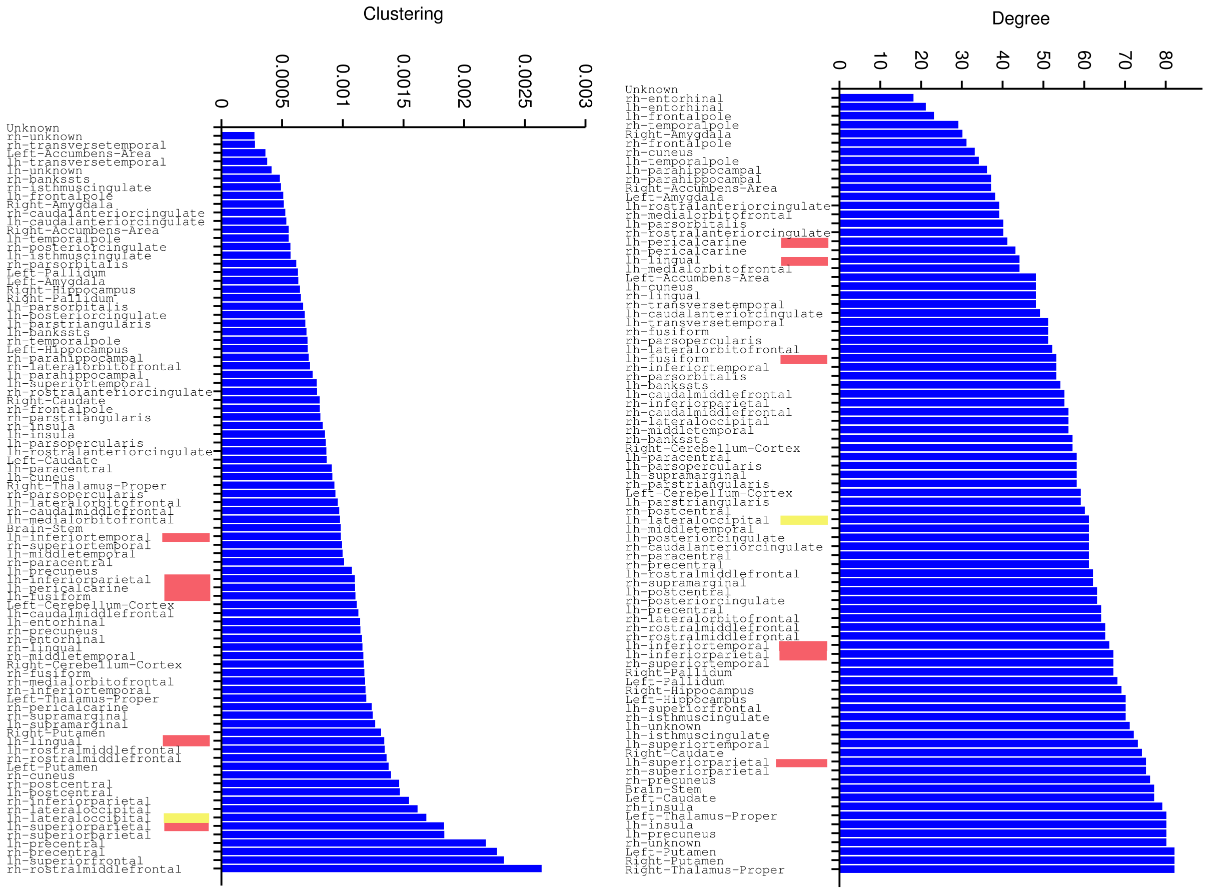


Figure 3: Left panel: Clustering value of each population. Right panel: Degree value of each population. Both properties are calculated on the structural connectivity matrix of the patient **CJ**. The clinical estimated EZ is highlighted in yellow, while the estimated PZ (predicted via Linear Stability Analysis) is highlighted in magenta.

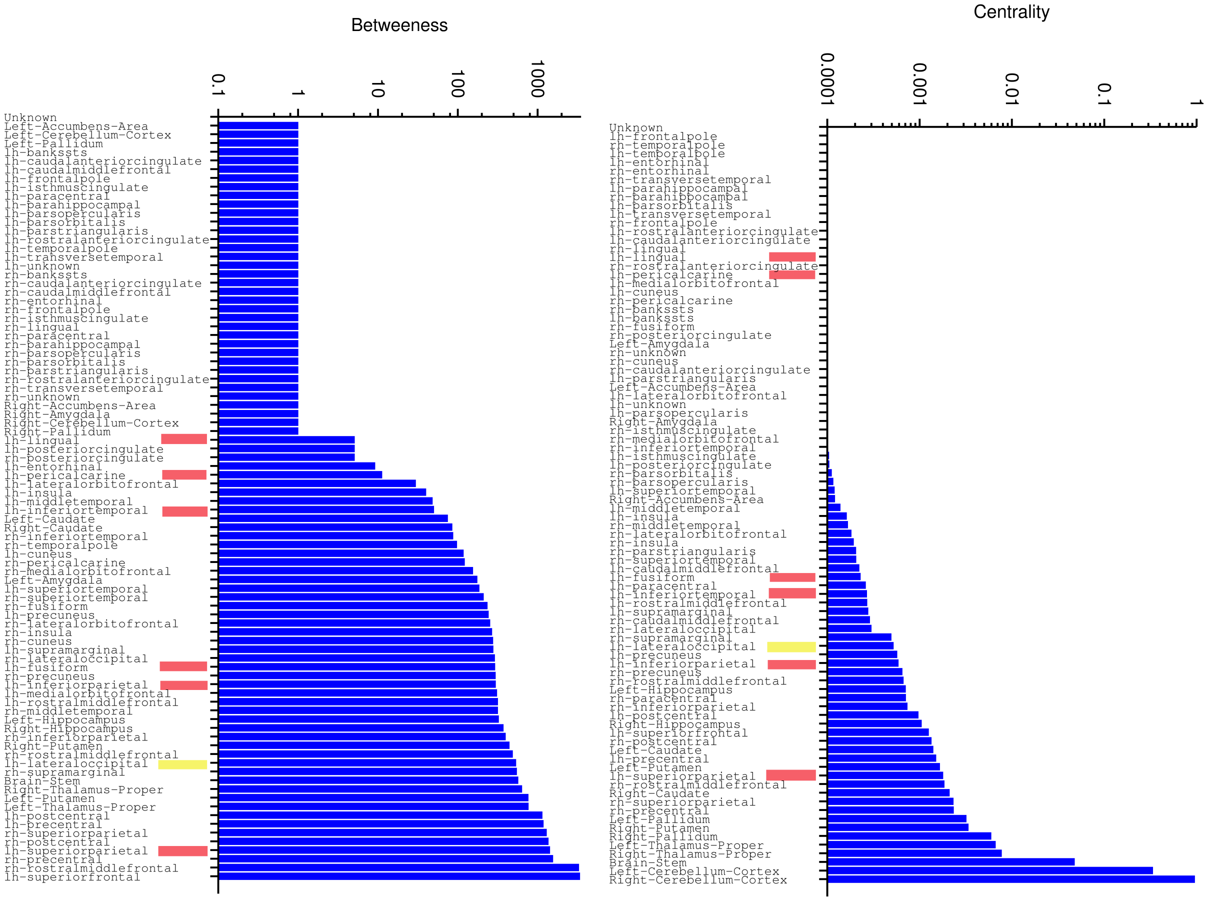


Figure 4: Left panel: Betweeness value of each population. Right panel: Centrality value of each population. Both properties are calculated on the structural connectivity matrix of the patient **CJ**. The clinical estimated EZ is highlighted in yellow, while the estimated PZ (predicted via Linear Stability Analysis) is highlighted in magenta.

### II. OTHER PATIENTS

Similar results to those shown in the main text has been obtained testing the procedure on the structural connectivity matrices of 15 patients that underwent a presurgical evaluation: this means that for all of them we knew the position of the epileptogenic zone thanks to clinicians' estimation (the clinical characteristics of each patient are given in Table **??**) and we could compare the predicted propagation zone with the one emerging from the analysis of SEEG data coming from implanted electrodes. However the analysis that we have done is not restricted to a particular matrix or epileptogenic zone; on the contrary is has a general applicability, as we have shown in Fig. 6(a) in the main text, and as we are going to show here for other six patients. Even though results depend on the location of the epileptogenic zone and on the subgraph outgoing from this zone, it is always possible to calculate the Linear Stability Analysis of the system, identify the propagation zone and the instability pathways and perform a minimal set of lesions on the connectivity network able to stop the seizure propagation.

*Patient FB*: FB shows Premotor epilepsy (right side). Engel score II. EZ: Precentral Gyrus. PZ prediction: Postcentral Gyrus, Caudal Middle Frontal Gyrus, Pars opercularis, Superior Frontal Gyrus, Thalamus, Putamen, Paracentral Cortex, Supramarginal Gyrus. PZ clinical prediction: Postcentral Gyrus, Superior Parietal Cortex. Lesions: it is necessary to cut all the links between the Precentral Gyrus and the nodes in the PZ (8) to stop the propagations (see Fig. **??**).

*Patient ET*: Parietal epilepsy in the left side. Engel score I. EZ regions: Posterior Cingulate Gyrus, Precuneus Cortex. PZ prediction: Isthmus-cingulate cortex, Postcentral Gyrus, Superior Parietal Cortex, Cuneus, Posterior Cingulate Gyrus, Parahippocampal Gyrus. PZ clinical prediction: Postcentral Gyrus, Superior Parietal Cortex. In this case both nodes of the EZ are contributing to destabilize the system, therefore it is necessary to remove the links between each epileptogenic zone and the propagation zones (11 links). In addition to this we have to look also at the propagation flow of the seizure and at the nodes that are immediately recruited due to the strength of their links, but that are not in the PZ (other 2 lesions) (see Fig. **??**).

*Patient AC*: Temporo-frontal epilepsy in the right side. Engel score III. EZ regions: Lateral Orbito Frontal Cortex, Temporal pole. PZ prediction: Rostral Middle Frontal Gyrus,

| Patient | Gender | Epilepsy duration (years) | Age at seizure onset (years) | Epilepsy type | Surgical procedure | Surgical outcome | MRI | Histopathology | Side |
|---|---|---|---|---|---|---|---|---|---|
| AC | F | 14 | 8 | Temporo-frontal | Sr | III | Anterior temporal necrosis | Gliosis | R |
| CJ | F | 14 | 9 | Occipital | Sr | III | N | FCD type1 | L |
| CM | M | 35 | 7 | Insular | GK | I | N | NA | L |
| CV | F | 18 | 5 | SMA | Sr | I | N | FCD type2 | L |
| ET | F | 23 | 7 | Parietal | Sr | I | FCD SPC | FCD type2 | L |
| FB | F | 16 | 7 | Premotor | Th | II | N | NA | R |
| FBO | M | 45 | 11 | Temporo-frontal | Sr | I | FCD F | FCD type2 | R |
| GC | M | 5 | 28 | Temporal | Sr | III | Temporopolar hypersignal | FCD type1 | R |
| IL | F | 18 | 20 | Occipital | N | NO | N | NA | R |
| JS | M | 11 | 18 | Frontal | Sr | I | Frontal necrosis (post-trauma) | Gliosis | R |
| ML | F | 10 | 17 | Temporal | Gk | II | Hippocampal sclerosis | NA | R |
| PC | M | 15 | 14 | Temporal | N | NO | N | NA | R |
| PG | M | 29 | 7 | Temporal | Sr | I | Cavernoma | Cavernoma | R |
| RB | M | 28 | 35 | Temporal | Sr | III | N | Gliosis | L |
| SF | F | 24 | 4 | Occipital | N | NO | PVH | NA | R |

Figure 5: Partial seizures were recorded with stereotactic EEG (SEEG) electrodes in 15 drug-resistant epileptic patients undergoing presurgical evaluation. The clinical characteristics of each patient are given in Table **??**: N, normal; L, left; R, right; Th, thermocoagulation; Gk, Gamma knife; Sr, surgical resection; NO, not operated; PVH, periventricular nodular heterotopia; FCD, focal cortical dysplasia; SPC, superior parietal cortex; F, Frontal; NA, not available.

Medial Orbito Frontal Cortex, Pars Orbitalis, Insula, Putamen, Pars Triangularis. PZ clinical prediction: Superior Frontal Gyrus, Rostral Middle Frontal Gyrus, (left) Lateral Orbito Frontal Cortex. In this case only one node of the EZ is contributing to the destabilization of the system and it is sufficient to cut the link between LOFC and RMFG to stop the propagation (see Fig. **??**).

*Patient GC*: Temporal epilepsy in the right side. Engel score III. EZ regions: Amygdala, Hippocampus. PZ prediction: Entorhinal Cortex, Parahippocampal Gyrus, Thalamus, Pallidum, Fusiform Gyrus. PZ clinical prediction: Superior Temporal Gyrus, Temporal Pole, Inferior Temporal Gyrus, Medial Orbito Frontal Cortex, Middle Temporal Gyrus, Parahippocampal Gyrus, Pars orbitalis, Pars triangularis, Rostral Middle Frontal Gyrus, Insula. Hippocampus plays a dominant role with respect to the Amygdala in the recruitment and propagation process, therefore it is sufficient to cut the links between this node and the nodes

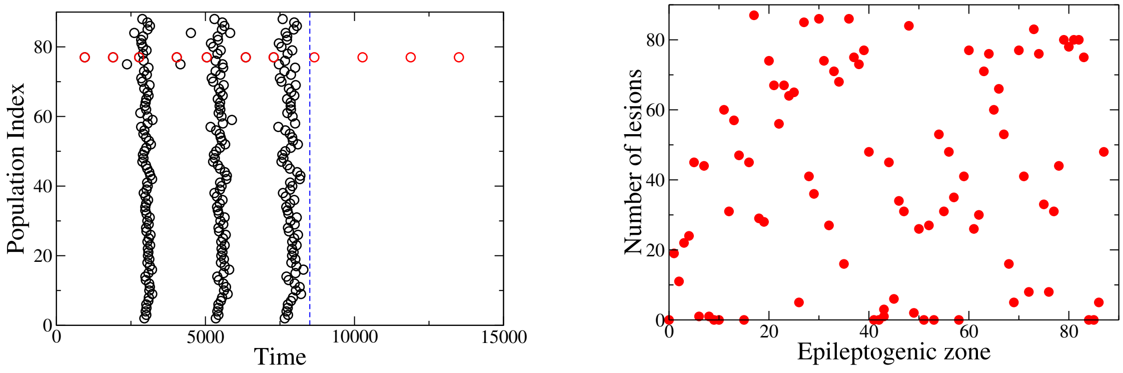


Figure 6: Analysis of patient **FB**. Left panel: Seizure events as a function of time. EZ (red dots): rh-Precentral Gyrus that corresponds to node 77. Disconnections PrG- PoG, PrG-CMFG, PrG-PoP, PrG-SFG, PrG-Th, PrG-Pu, PrG-PaC, PrG-SMG must be performed in order to stop the seizure. All the disconnections are performed among populations belonging to the right hemisphere. Dashed blue line corresponds to the time at which lesions occur. Right panel: Number of lesions necessary to stop seizure propagation as a function of the epileptogenic region.

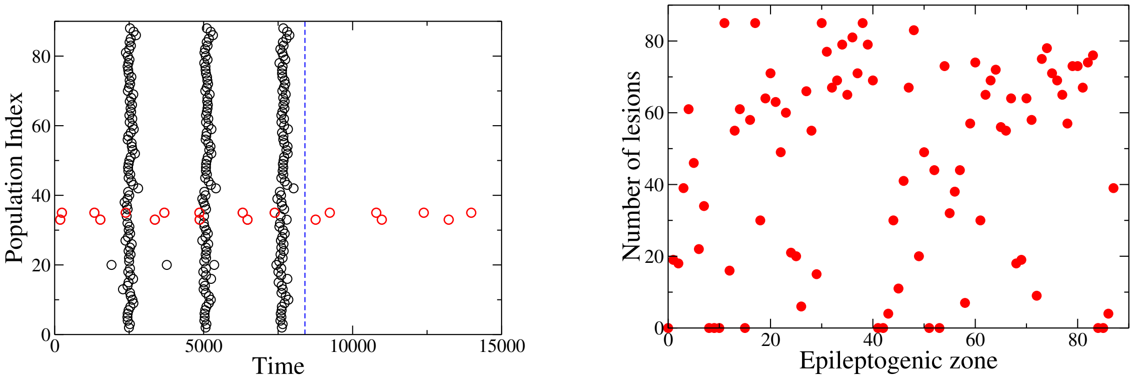


Figure 7: Analysis of patient **ET**. Left panel: Seizure events as a function of time. EZ (red dots): lh-Posterior Cingulate Gyrus, lh-Precuneus Cortex corresponding respectively to nodes 33 and 35. Partial targeted disconnection procedure to be performed in order to stop seizure propagation: PCunC-SPC, PCunC-PCG, PCunC-ICC, PCunC-Cun, PCunC-rPCunC, PCunC-Pac, and the links between PCG and nodes SPC, ICC, Cun, rPCunC, Pac, SFG, CACC. The targeted disconnection is performed among populations in the left hemisphere. Dashed blue line corresponds to the time at which lesions occur. Right panel: Number of lesions necessary to stop seizure propagation as a function of the epileptogenic region.

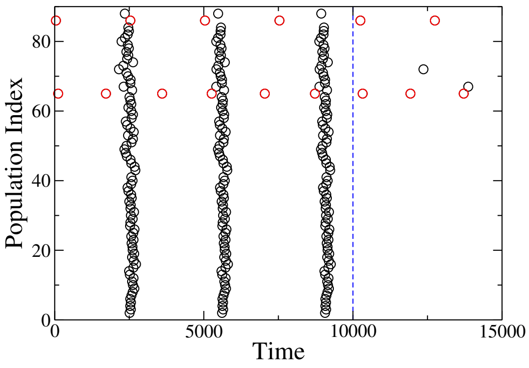


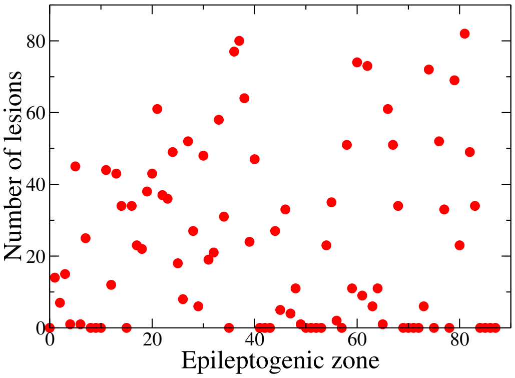


Figure 8: Analysis of patient **AC**. Left panel: Seizure events as a function of time. EZ (red dots): rh-Lateral Orbito Frontal Cortex, rh-Temporal pole corresponding respectively to nodes 65 and 86. Partial targeted disconnection procedure to be performed in order to stop seizure propagation: LOFC-RMFG. The targeted disconnection is performed among populations in the right hemisphere. Dashed blue line corresponds to the time at which lesions occur. Right panel: Number of lesions necessary to stop seizure propagation as a function of the epileptogenic region.

belonging to the PZ (5 links in total), in addition to the links between Amigdala and Hippocampus, Amigdala and Temporal Pole to stop the seizure. The former link (Amg-TmP) is very strong and the Temporal Pole turns out to be the biggest element of the maximal eigenvector after the nodes belonging to the PZ (see Fig. **??**).

*Patient CV*: CV shows Supplementary Motor Areas epilesy (left side). Engel score I. EZ regions: Posterior Cingulate Gyrus, Caudal Middle Frontal Gyrus, Superior Frontal Gyrus. PZ prediction: Precentral Gyrus, Caudal Middle Frontal Gyrus, Rostral Middle Frontal Gyrus, (right) Superior Frontal Gyrus, Caudal Anterior Cingulate Cortex, Paracentral Cortex. PZ clinical prediction: Precentral Gyrus, Postcentral Gyrus. It turns out that 1 out of 3 areas in the EZ has a higher impact on the system instability, thus being determinant to the recruitment process. When more that one area constitutes the EZ it is necessary to take into account also the links among the areas, which sustain and enhance the propagation, as testified in this case by the fact that it is necessary to disconnect the link PCG-CMFG to achieve the desired goal. In total, it is sufficient to perform 3 lesions in order to stop the seizure propagation (see Fig. **??**).

*Patient RB*: Temporal epilepsy in the left side. Engel score III. EZ regions: Amygdala, Hippocampus, Entorhinal cortex, Fusiform gyrus, Temporal Pole, right Entorhinal cortex.

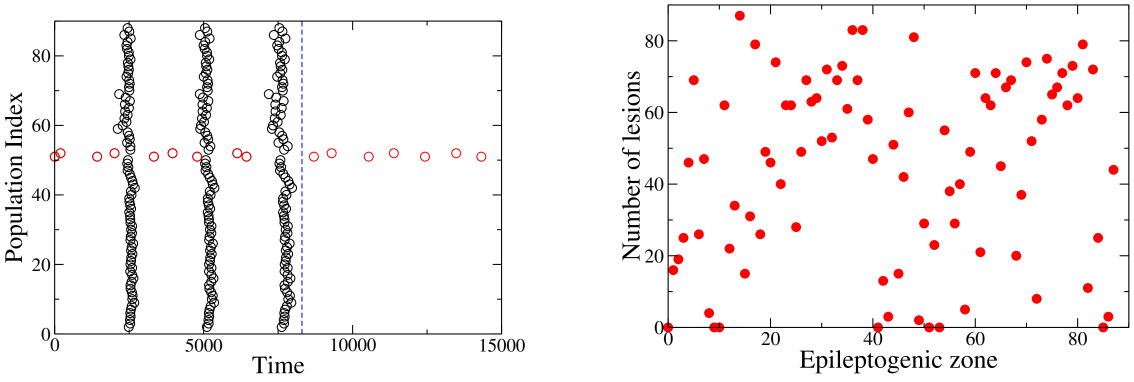


Figure 9: Analysis of patient **GC**. Left panel: Seizure events as a function of time. EZ (red dots): rh-Hippocampus, rh-Amygdala corresponding respectively to nodes 51 and 52. Partial targeted disconnection procedure to be performed in order to stop seizure propagation: Hi-EntC, Hi-PHiG, Hi-Pal, Hi-Th, Hi-FuG, Hi-Amg, Amg-TmP . The targeted disconnection is performed among populations in the right hemisphere. Dashed blue line corresponds to the time at which lesions occur. Right panel: Number of lesions necessary to stop seizure propagation as a function of the epileptogenic region.

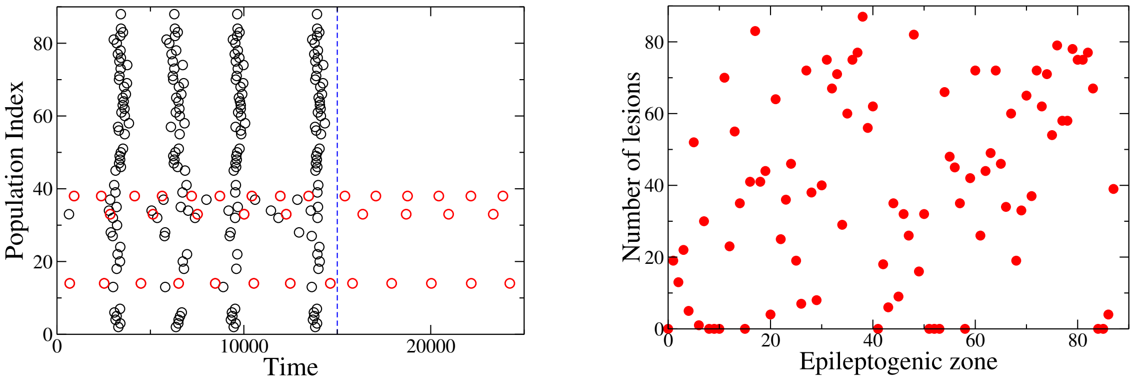


Figure 10: Analysis of patient **CV**. Left panel: Seizure events as a function of time. EZ regions (red dots): Posterior Cingulate Gyrus (node 33), Caudal Middle Frontal Gyrus (node 14), Superior Frontal Gyrus (node 38). All the areas belonging to the EZ are located in the left hemisphere. Links between regions SFG-RMFG, SFG-(rh-SFG) and PCG-CMFG are cut in order to stop the seizure. Dashed blue line corresponds to the time at which lesions occur. Right panel: Number of lesions necessary to stop seizure propagation as a function of the epileptogenic region.

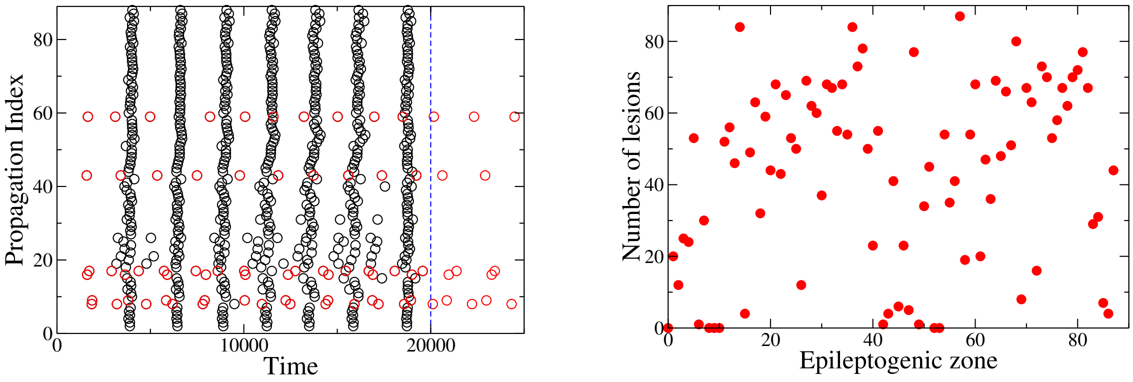


Figure 11: Analysis of patient **RB**. Left panel: Seizure events as a function of time. EZ regions (red dots): Amygdala (node 9), Hippocampus (node 8), Entorhinal cortex (node 16), Fusiform gyrus (node 17), Temporal Pole (node 43), rh-Entorhinal cortex (node 59). All the areas belonging to the EZ are located in the left hemisphere. Links between all the regions in the EZ and the regions in the PZ must be cut in order to stop the seizure. Dashed blue line corresponds to the time at which lesions occur. Right panel: Number of lesions necessary to stop seizure propagation as a function of the epileptogenic region.

PZ prediction: Inferior Temporal Gyrus, Lateral Occipital Cortex, Lingual gyrus, Parahippocampal Gyrus, Cerebellum Cortex. PZ clinical prediction: Superior Temporal Gyrus, Middle Temporal Gyrus, Inferior Temporal Gyrus, Insula, Parahippocampal Gyrus. Links between all the regions in the EZ and the regions in the PZ must be cut in order to stop the seizure (see Fig. **??**).

---


[1] Jirsa, V. K., Stacey, W. C., Quilichini, P. P., Ivanov, A. I., Bernard C. *On the nature of seizure dynamics*, Brain 137:2210 -2230 (2014).

[2] Haken, H. *Advanced synergetics: Instability hierarchies of self-organizing systems and devices* (Vol. 20). Springer Science & Business Media (2012).

[3] Naze, S., Bernard, C. and Jirsa, V., *Computational modeling of seizure dynamics using coupled neuronal networks: factors shaping epileptiform activity*, PLoS computational biology 11(5), e1004209 (2015).

[4] Pohjanpalo, H., *System identifiability based on the power series expansion of the solution*,

Math. Biosci. 41, 21-33 (1978); Lecourtier, Y. et al., *Volterra and generating power series approaches to identifiability testing.* In Identifiability of Parametric Models. Pergamon Press, Oxford (1987); Vajda, S. et al., *Similarity transformation approach to identifiability analysis of nonlinear compartmental models*, Math. Biosci. 93, 217-248 (1989); Ljung,L. and Glad, T., *On global identifiability for arbitrary model parametrizations*, Automatica 30, 265-265 (1994); Margaria, G. et al., *Differential algebra methods for the study of the structural identifiability of rational function state-space models in the biosciences*, Math. Biosci. 174, 1-26 (2001); White, L. et al., *The structural identifiability and parameter estimation of a multispecies model for the transmission of mastitis in dairy cows*, Math. Biosci. 174, 77-90 (2001).

[5] Proix, T., Bartolomei, F., Chauvel, P., Bernard, C., Jirsa, V. K. *Permittivity Coupling across Brain Regions Determines Seizure Recruitment in Partial Epilepsy*, J. Neurosci. 34, 15009-15021 (2014).

[6] Arnold, V., Afraimovich, V., Ilyashenko, Y., Shilnikov, L. *Bifurcation theory.* In: Arnold, V. (ed.) Dynamical Systems. Encyclopaedia of Mathematical Sciences, vol. V. Springer, Berlin (1994); Mischenko, E., Kolesov, Y., Kolesov, A., Rozov, N. *Asymptotic Methods in Singularly Perturbed Systems. Monographs in Contemporary Mathematics.* Consultants Bureau, New York (1994).

[7] Proix, T., Bartolomei, F., Guye, M. and Jirsa, V.K. *Individual structural connectivity defines propagation networks in partial epilepsy*, Brain 140:641-654 (2017).

[8] Shimada, I., and Nagashima, T. *A numerical approach to ergodic problem of dissipative dynamical systems*, Progress of Theoretical Physics 61.6: 1605-1616 (1979). Benettin, G., et al. *Lyapunov characteristic exponents for smooth dynamical systems and for Hamiltonian systems; a method for computing all of them. Part 1: Theory*, Meccanica 15.1: 9-20 (1980).
